# Nucleus Basalis Stimulation Enhances Working Memory by Stabilizing Attractor Networks in Prefrontal Cortex

**DOI:** 10.1101/674465

**Authors:** Xue-Lian Qi, Ruifeng Liu, Balbir Singh, David Bestue, Albert Compte, Almira I. Vazdarjanova, David T. Blake, Christos Constantinidis

## Abstract

Acetylcholine in the neocortex is critical for executive function. Degeneration of the basal forebrain cholinergic system is associated with cognitive decline in aging and Alzheimer’s disease. Cholinergic agonists and acetylcholinesterase inhibitors improve cognitive performance as does intermittent electrical stimulation of the cortical source of acetylcholine, the Nucleus Basalis (NB) of Meynert. Here we tested how NB stimulation improves working memory behavior and alters its neural code. NB stimulation increased dorsolateral prefrontal activity during the delay period of working memory tasks but did not strengthen phasic responses to the optimal visual stimulus of each neuron. Unexpectedly, improvement of behavioral performance was not the result of increased neural selectivity. Tuning of neuronal responses broadened, which rendered an attractor network more stable and filtered distracting visual stimuli more effectively. Thus, the effects of acetylcholine on prefrontal neural activity and selectivity in working memory contrast those of dopamine and stabilize neural ensembles based on neuromodulatory tone.

## Introduction

The forebrain cholinergic system tightly regulates higher cognitive function (Galvin et al., 2018; Sarter et al., 2016). Losses in cognitive performance with aging and Alzheimer’s disease occur in parallel with degeneration of the brain’s cholinergic systems, and cholinergic deficits correlate tightly with the degree of cognitive decline (Ballinger et al., 2016; Terry and Buccafusco, 2003). Cholinesterase inhibitors, which prolong the neurotransmitter’s ability to stimulate post-synaptic receptors and amplify the natural pattern of acetylcholine release, are frontline medications for treating Alzheimer’s disease (Birks and Harvey, 2018).

Improvement in cognitive functions may alternatively be achieved by stimulation of the Nucleus Basalis (NB) of Meynert (Freund et al., 2009), the sole source of neocortical acetylcholine in humans and other primates (Mesulam et al., 1983). This approach offers distinct benefits over drug administration, including avoidance of peripheral cholinergic stimulation; optimized timing; and activation of non-cholinergic projection neurons also found in Nucleus Basalis, which are normally co-active with the cholinergic fibers (Kim et al., 2015; Walker et al., 1989). Stimulation of basal forebrain neurons has been shown to be effective in the context of neuroplasticity (Bakin and Weinberger, 1996; Brzosko et al., 2019; Froemke, 2015; Kilgard and Merzenich, 1998), and in enhancing behavioral performance of attention and memory tasks at levels comparable to high doses of acetylcholinesterase inhibitors, with improvements that aggregate rather than attenuate over time (Blake et al., 2017; Liu et al., 2018, 2017).

We were therefore motivated to examine how intermittent NB stimulation affects neuronal activity so as to improve performance of working memory tasks. We focused specifically on the prefrontal cortex, an area critical for working memory and cognitive plasticity (Constantinidis and Klingberg, 2016), which receives innervation from a dedicated sub-region of the Nucleus Basalis (Gielow and Zaborszky, 2017). Pharmacological studies suggest that iontophoresis of cholinergic agonists into the prefrontal cortex increases neuronal activity specifically for the neuron’s preferred stimulus, therefore sharpening its neuronal tuning (Dasilva et al., 2019; Sun et al., 2017; Yang et al., 2013), whereas inhibitors depress prefrontal activity (Major et al., 2015; Yang et al., 2013). Despite our expectations based on these results, widespread cholinergic modulation through NB stimulation revealed no phasic increases in prefrontal activity during the preferred visual stimulus presentation, and broadening rather than narrowing of visual stimulus selectivity. Our results lead to a reevaluation of mechanisms through which neuromodulatory tone improves cognitive function, in the context of attractor models of working memory generation (Compte et al., 2000; Wimmer et al., 2014) and efficient coding models (Ma et al., 2006; Zhang and Sejnowski, 1999).

## Results

Two monkeys were implanted with unilateral electrodes targeting the Nucleus Basalis to determine the effects of stimulation on behavioral performance and neural activity. Electrode placement was guided by MR imaging (Fig. 1A). To obtain functional confirmation of the targeting, we collected LFP signals from the implanted electrode with the monkey at rest, with and without NB stimulation (Bjordahl et al., 1998). Continuous NB stimulation at 80 Hz produced LFP desynchronization (Fig. 1B, and S1). Power in the 5-15 Hz range was significantly lower during NB stimulation than control (paired t-test, t_17_=3.14, p=0.006 and t_33_=2.5, p=0.02 for the two subjects, respectively). Electrode placement was confirmed with ChAT immunohistochemistry, post mortem (Fig. 1C-E). Control sections where the primary antibody was omitted produced no distinct cell-specific staining.

**Figure 1.**
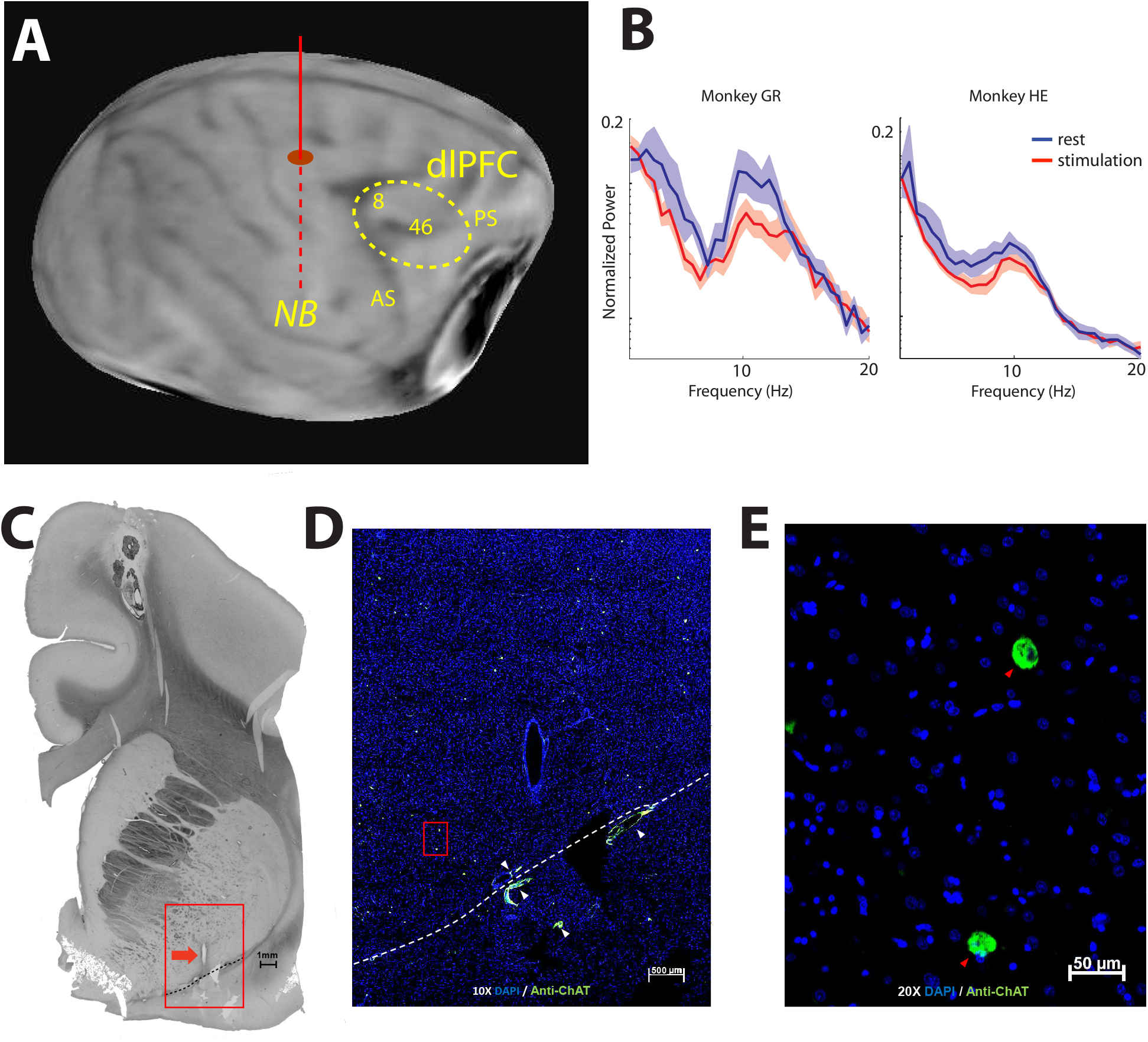
Localization and effects of stimulation. **(A)** Anatomical MR scan from one monkey obtained prior to implantation. The approximate location of the implanted electrode is indicated with the solid/dashed vertical line. The dotted area represents the cortical region sampled with neurophysiological recordings. Abbreviations, AS: arcuate sulcus; PS: principal sulcus. (**B)** Power Spectrum of Local Field Potential recorded from the implanted electrode during rest and following 80 Hz stimulation in the two animals (subjects HE and GR). **(C)** Histology and ChAT Immunohistochemistry. A 50 micron thick coronal plane section is viewed in light microscopy displaying the most inferior track of the electrode cannula. The red box surrounds the cannula track (red arrow) and indicates the area enlarged in section D. The dashed line marks the floor of the Nucleus Basalis. **(D)** The marked area enlarged shown under merged green and blue fluorescence. Blue color marks nuclei with DAPI, and the green marks antibody to ChAT. White arrows mark blood vessel autofluorescence. The dashed line is the same as in A and marks the floor of the basal forebrain. **(E)** Nucleus Basalis at 20X magnification. DAPI (blue), Anti-ChAT (green), Anti-ChAT-containing neurons (red arrows), Blood vessel autofluorescence (white arrows).

### Behavioral performance

The monkeys performed a task requiring them to view two stimuli presented in sequence and make an eye movement to the remembered location of either the first or second of them (Fig. 2A). A white fixation point instructed the monkeys to remember the first stimulus; a blue fixation point required them to remember the second stimulus. The two stimuli could appear at any of eight possible locations, presented in a randomized order (see insets in Fig. S2 and S3). When NB stimulation was delivered, it was applied during an inter-trial interval between two trials for 15 s, at a frequency of 80 pulses per s (Fig. 2B-D). The monkeys performed consecutive trials for 45 s (typically 4-5 completed trials), without NB stimulation. At the end of the trial that exceed the 45 s threshold, NB stimulation was applied anew and the cycle was repeated.

**Figure 2.**
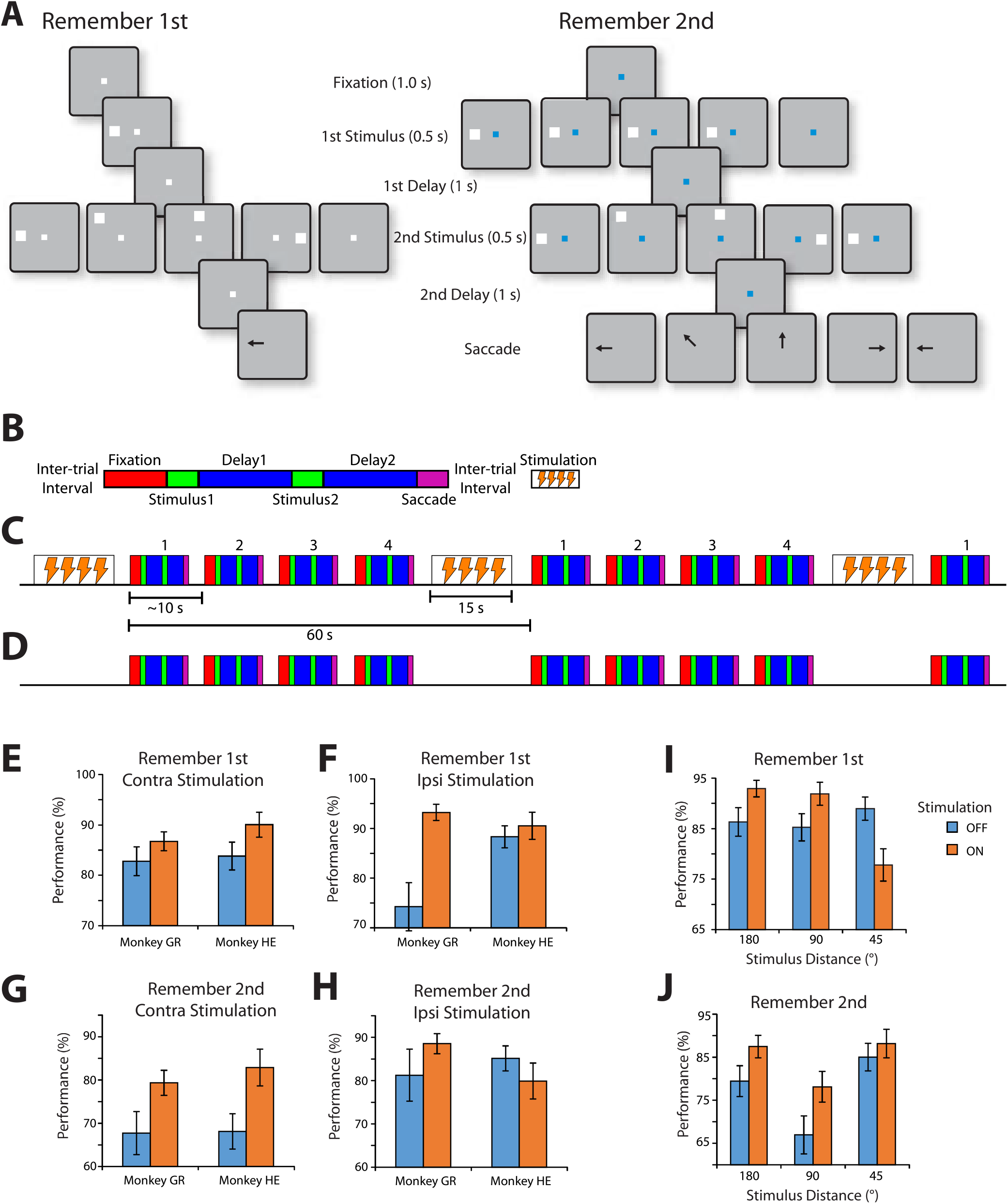
Behavioral task and performance. **(A)** Successive frames illustrate the sequence of events in the behavioral task. Depending on the white or blue color of the fixation point, the monkey has to remember either the first or the second of two visual stimuli presented in sequence, respectively. At the end of the trial, the fixation point turns off and the monkey needs to perform an eye movement towards the remembered location of the visual stimulus in order to receive a liquid reward. **(B).** Schematic diagram of a single trial of the task. Blocks represent the successive events in the task: fixation, first visual stimulus presentation, first delay period, second visual stimulus presentation, second delay period, and saccade. Successive trials are separated by intertrial intervals. NB stimulation, when delivered, always occurs during the intertrial interval. **(C)** Sequence of trials during a stimulation block, in a compressed time scale, relative to panel B. Successive trials, labeled 1-4, each lasting approximately 10 seconds, are followed by 15 s of stimulation. The precise duration of a trial differed depending on how quickly the animal initiated the trial, and if the trial was successfully completed, or aborted, e.g. due to a break in fixation. **(D)** Sequence of trials during a control (no-stimulation) block. The trials are arranged exactly in the same fashion as in the stimulation block, including an extended intertrial interval every 60 s, during which however no stimulation is applied (sham). (**E-H),** Percentage of correct trials is shown for each of the two monkeys, for different visual stimulus types (n=18 sessions for stimulation, 17 for control for monkey GR; n=19 stimulation and 35 control for monkey HE). **(E)** Mean performance (and sem) for trials in which first visual stimulus appears contralateral to the stimulation site, when the monkey is executing the remember-first task, and needs to remember the first visual stimulus. **(F)** Performance in the remember-first task when the first visual stimulus appears ipsilateral to the stimulation site. **(G)** Performance in the remember-second task, when the second visual stimulus appears contralateral to the stimulation site. **(H)** Performance in the remember-second task when the second visual stimulus appears ipsilateral to the stimulation site. **(I)** Performance in the remember-first task, for trials grouped by distance between the first and second visual stimulus (180, 90, or 45°), under stimulation or control conditions. Data from both monkeys pooled together. (**J)** Performance in the remember-second task, for trials grouped by distance.

NB stimulation was delivered in two modes. In the first mode, daily sessions were performed without any NB stimulation and these were preceded or followed by daily sessions during which intermittent NB stimulation was performed based on the schedule indicated in Fig. 2C. These daily sessions allowed us to collect a large number of trials and characterize the behavioral performance of the monkeys (Fig. 2E-H). In the second mode, when neuronal recordings commenced, the monkeys performed blocks of 60 trials without NB stimulation, followed by blocks of trials with NB stimulation, which in this case, too, was applied in intermittent fashion, during the inter-trial interval of stimulation blocks.

Behavioral performance improved with intermittent NB stimulation delivered in daily sessions (Fig. 2E-H). For monkey GR, improvement was observed for all conditions. A 3-way ANOVA on performance with factors a) NB stimulation (on or off), b) task (remember-first or remember-second), and c) location of visual stimuli (contralateral or ipsilateral visual stimulus to be remembered) revealed a significant main effect of NB stimulation (*F*_*1,128*_=19.6, p=2.06×10^−5^). For monkey HE, stimulation improved performance specifically when the visual stimulus to be remembered was at a location contralateral to the site of the stimulation electrode (Fig. 2E-H). NB Stimulation was ineffective when the visual stimulus was ipsilateral to the stimulation (Fig. 2F,H). Performing the same 3-way ANOVA analysis revealed no net effect of NB stimulation, precisely because of the opposing effects in the two hemifields (*F*_*1,204*_=0.27, p=0.6), but now a significant three-way interaction between task, stimulation, and side of visual stimulus (*F*_*1,128*_=6.04, p=0.015). Considering the contralateral conditions alone (Fig. 2E-G) the effect of NB stimulation was highly significant (3-way ANOVA with factors monkey, and stimulation: *F*_*1,173*_ = 14.14, p= 0.0002 for remember-first task; *F*_*1,173*_= 5.61 p=0.02 for remember-second task). Dissecting the effect of stimulation for different combinations of stimulus locations additionally revealed that NB stimulation decreased performance for visual stimuli appearing at adjacent locations (Fig. 2I, 45° distance), with excess errors directed to the location of the distractor (two-tailed t-test, *t*_*66*_=2.83, p=0.006). This pattern of responses was not observed for NB stimulation in the remember-second condition, for a second visual stimulus appearing at a close distance to the initial distractor (45° condition in Fig. 2J). Modeling results provided an explanation for this pattern of behavioral effects (discussed below). In summary, NB stimulation produced an overall improvement in performance, however the effect depended on the relative position of stimuli relative to the site of stimulation and distractors.

### Overall effects on neural activity

We recorded from a total of 233 neurons (102 and 131 in the two monkeys, respectively) in areas 8 and 46 of the dorsolateral prefrontal cortex with and without Nucleus Basalis stimulation, always delivered in the inter-trial interval. Recording cylinders were implanted on the same side as the stimulation electrode. NB stimulation had a predominantly excitatory effect in neural activity recorded in correct trials (Fig. 3A-D). Mean firing rate averaged across all neurons (Fig. 3E) during the fixation period was 10.9 sp/s in control trials and 13.4 sp/s in NB stimulation trials, a significant difference (two-tailed paired t-test, *t*_*232*_=4.21, p=3.65×10^−5^). A similar trend was observed in both monkeys (10.4 vs 11.9 sp/s, p=1.4×10^−4^; 11.3 vs. 14.6 sp/s, p=0.07 for subjects GR and HE, respectively). During the visual stimulus presentation period, mean firing rate (averaged across all visual stimulus conditions and neurons) was 13.4 sp/s for the control and 16.2 sp/s for the NB stimulation condition (two-tailed paired t-test, *t*_*232*_=4.39, p=1.7×10^−5^). A similar trend was observed in both monkeys (13.4 vs 16.0 sp/s, p=9.5×10^−5^; 13.3 vs. 16.3 sp/s, p=0.023 for the two animals, respectively). However, the increase in firing rate was not uniform across stimuli, and was accounted mostly by increased responsiveness to suboptimal stimuli, rather than the best stimulus of each neuron, which remained relatively unchanged (discussed further, below).

**Figure 3.**
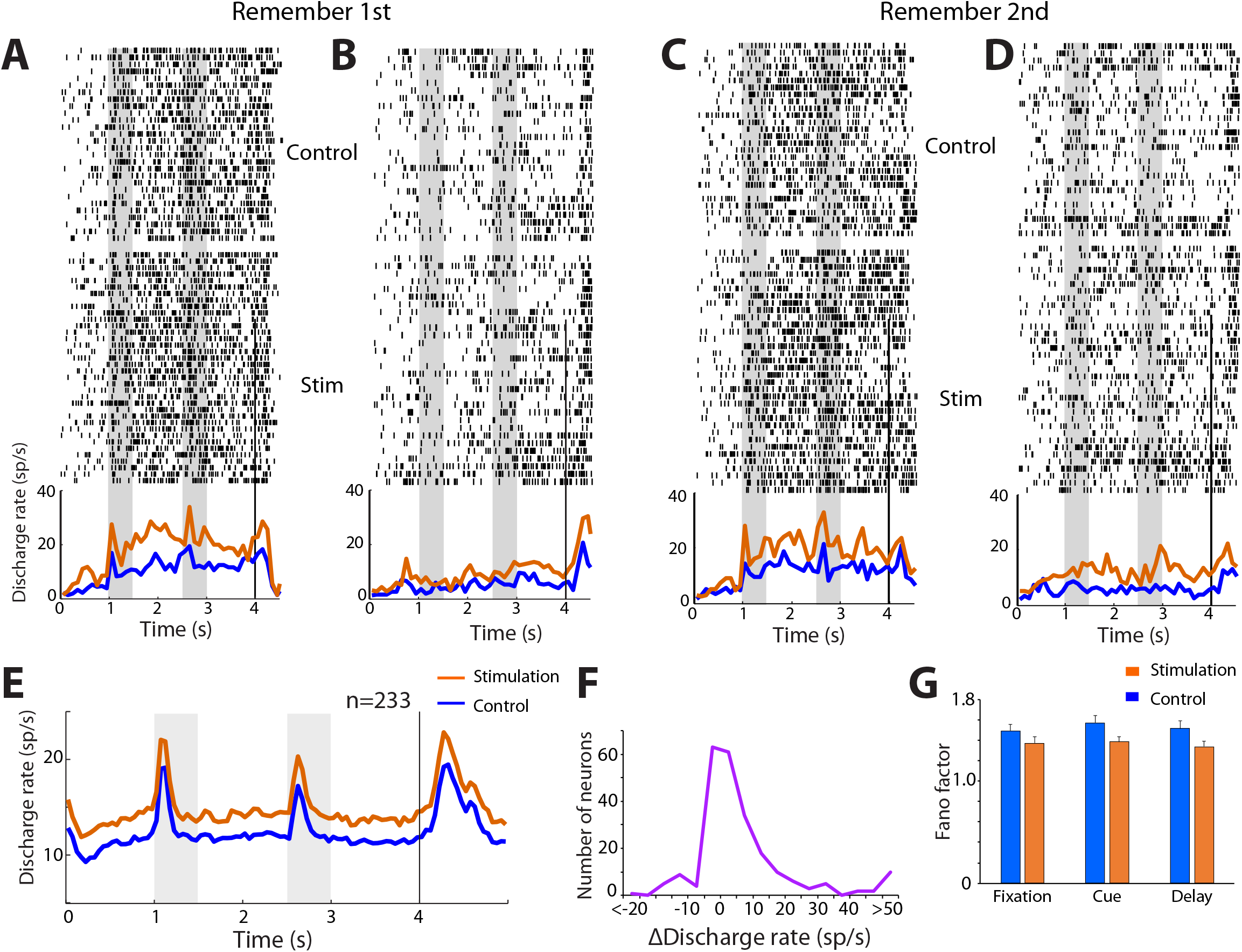
Neuronal stimulation effects. (**A-D)**. Raster plots represent responses of a single prefrontal neuron in the remember-first (A-B) and remember-second task (C-D), under control and NB stimulation conditions. Line traces represent averaged, peri-stimulus time histograms, with and without NB stimulation. Trials are pooled from conditions when the first visual stimulus appeared to the contralateral (A, C) or ipsilateral hemifield (B, D). Gray bars indicate time of visual stimulus presentation. Line traces represent averaged, peri-stimulus time histograms, with and without stimulation. (**E)**. Population PSTH averaged across all neurons (n=233) and all stimulus conditions. (**F)**. Distribution of changes in firing rate in the fixation period across all neurons (n=233). (**G)**. Mean Fano factor of spike counts averaged over the fixation, cue, and first delay period of the task (n=233).

The effects of NB stimulation were not uniform across neurons, either (Fig. 3F). NB stimulation produced a significant increase in fixation period firing rate for 97/233 neurons (estimated with a t-test, at the 0.05 significance level), no significant difference in 99/233 neurons, but also a decrease in firing rate for 37/233 of neurons. The distribution of firing rate differences computed in blocks of trials with or without NB stimulation deviated significantly from a normal distribution (Fig. 3F, KS test for normalized rate differences, compared to normal distribution, p=8.74×10^−6^).

NB stimulation did not only affect firing rates, but also their variability. A small but significant decrease in Fano factor, a measure of trial to trial variability, was observed under NB stimulation (Fig. 3G). The reduction over the control condition was evident in all task intervals. A 2-way ANOVA, with factors task-interval and stimulation showed a significant main effect of stimulation (*F*_*1,1353*_ =10.1, p=0.0015) but no interaction (*F*_*2,1353*_=0.17, p=0.8).

### Effects in different task intervals

Among neurons recorded, 112 (67 and 45 from the two subjects respectively) responded to the visual stimuli (evaluated with a paired t-test, at the p<0.05 significance level during the visual stimulus presentation or delay period). We examined these neurons in more detail, as they were revealing of the effects of NB stimulation in the context of the task. For a total of 54 neurons that responded to visual stimuli (41 and 13 from subjects GR and HE, respectively), firing rate in the fixation period was significantly higher after NB stimulation (evaluated with a t-test at the p<0.05 level). An example is shown in Fig. 3A-D. For 16 neurons, firing rate was significantly decreased. We examine these populations of neurons separately as exemplars demonstrating the range of changes in activity, below, and in the Supplemental Information.

For the neurons with an overall increase in activation during the fixation interval, the effect was stable over the time course of ~1 min between cycles of repeated stimulation (see Fig. 2B-C). The firing rate was elevated in the first trial following stimulation and no decrease was evident in the 4-5 successive trials that followed until the next train of stimulation was applied (Fig. 4H, red line). Interestingly, NB stimulation did not produce uniform effects across all task intervals. In the same group of neurons, NB stimulation had no effect during the inter-trial interval (Fig. 4A, 2-way ANOVA with factors tasks and stimulation: main effect of stimulation *F*_*1,53*_ =2.6, p=0.113). The phasic response to the preferred visual stimulus itself (peak in ~200 ms after visual stimulus) was also largely unchanged between the control and NB stimulation conditions (Fig. 4A, 2-way ANOVA with factors tasks and stimulation: main effect of stimulation *F*_*1,53*_ =0.397; p=0.53). The absence of an enhancement to the preferred visual stimulus response was evident in both the remember-first and remember-second tasks (Fig. S2, S3), and for both the first and second presentation of a visual stimulus in the receptive field (Fig. 4A). On the other hand, the increase in firing rate in trials with NB stimulation persisted during the delay periods after each stimulus presentation (2-way ANOVA with factors tasks and stimulation: main effect of stimulation *F*_*1,53*_ =6.20; p=0.016 for first delay period, *F*_*1,53*_ =6.42; p=0.014 for second delay period). These results of NB stimulation moved in the same direction for both remember-first and remember-second tasks (Fig. S4A), and for both monkeys (Fig. S4B-C).

**Figure 4.**
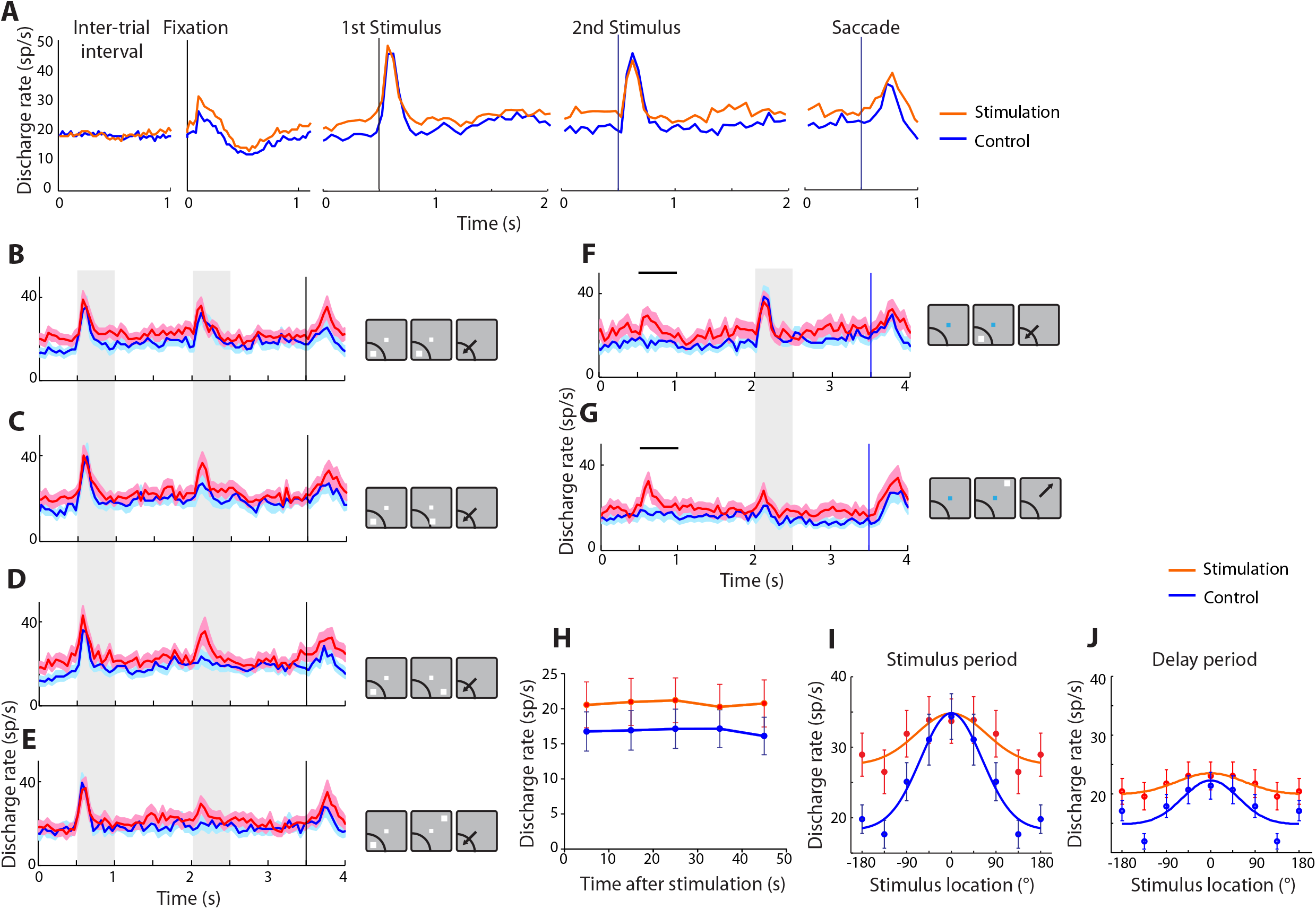
Population responses under stimulation. (**A)** Mean firing rate with and without NB stimulation is shown during the intertrial interval, fixation interval, first visual stimulus presentation involving the best visual stimulus of each neuron, second visual stimulus presentation involving the best visual stimulus of each neuron, and saccade towards best visual stimulus. Results from the remember-second task are shown, for neurons with significant increase in activation by NB stimulation (n=54 neurons for all panels). (**B-E),** Mean firing rate in the remember-first task, in conditions involving presentation of the first visual stimulus in the receptive field, followed by a second visual stimulus at progressively less responsive locations. Gray bars represent the times of visual stimulus presentations. Insets to the right of PSTH represent location of the visual stimuli relative to each neuron’s receptive field; the receptive field is shown always at the same location for simplicity; results from neurons with different receptive field locations have been averaged together. **(F-G),** Mean firing rate in the remember-second task, in conditions involving no first visual stimulus, followed by a second visual stimulus in or out of the receptive field. Horizontal lines illustrate the times that a first visual stimulus would have been delivered relative to the onset of the fixation point, had one been present in these trials. **(H)** Firing rate in sequential trials in blocks of trials when stimulation was applied or not. Abscissa represents time after the offset of stimulation, or sham inter-trial interval. **(I)** Population tuning curve for the cue period, obtaining by averaging responses of individual neurons to visual stimuli relative to each neuron’s preferred reference location (depicted at 0°). **(J)**. As in I, for the delay period.

### Neural tuning

Although NB stimulation did not improve responses for the best visual stimulus location, it enhanced responses to visual stimuli at non-optimal locations, which resulted in broadening of receptive fields during the cue presentation and delay period (second visual stimulus in Fig. 4C-E; first visual stimulus in Fig. S2F-J). The broadening of the receptive fields was also evident in the remember-second task (Fig. S3F-J). The population tuning curve based on the second-visual stimulus location illustrated the effect (Fig. 4I). In order to quantify differences in responsiveness to sub-optimal visual stimuli, we relied on a selectivity index (SI) defined as (Max-Min)/(Max+Min) where Max and Min represent the firing rate to the best and worst stimulus location for each neuron. The NB stimulation condition produced a significantly lower SI value, in both monkeys (2-way ANOVA with factors remember-first/remember-second tasks and stimulation: main effect of stimulation *F*_*1,53*_ = 25.4, p = 5.7×10^−6^ for cue period, and *F*_*1,53*_ = 24.2, p = 8.7×10^−6^ for delay period - Fig. S5A-D). Unlike the decrease in selectivity for visual stimulus location in prefrontal neuronal activity, the representation of task information (higher firing rate for remember-first or remember-second) was relatively unaffected by NB stimulation. We quantified this change by relying on 2-way ANOVA analysis with factors task (remember-first or remember-second) and stimulation (on or off), plotting the p-value for the main effect of task on firing rate (Fig. S6A-B). Effects of NB stimulation in the delay period firing rate following the first and second visual stimulus in the remember-first and remember-second task can also be seen in Fig. S6C-F.

### Bump Attractor Model

The decrease in spatial selectivity of individual neurons we observed under NB stimulation results in more neurons across the population being excited after the presentation of any visual stimulus. If we consider the activity of excited neurons forming a bump of activity in the network, then effects of NB stimulation can be conceptualized as broadening of this bump in the population of prefrontal neurons, which is thought to act as an attractor network during working memory (Compte et al., 2000; Inagaki et al., 2019). A bump attractor model that relies on excitation between similarly tuned neurons to maintain stimulus-selective, persistent spiking even after the stimulus is no longer present, captures many of the properties of working memory behavior (Barbosa et al., 2020; Wimmer et al., 2014). The model makes interesting predictions for behavioral performance under different combinations of remembered visual stimulus and distractor, of remember-first and remember-second conditions (Compte et al., 2000), and under bump broadening elicited by a general increase in network excitability (simulating the NB stimulation “ON” condition; see also Supplemental Information). A broader bump leads to a more stable bump attractor, less sensitive to noise fluctuations in the network (Fig. 5E,F,G,H), thus leading to more accurate read-outs at the end of the delay (Fig. 5J,K,M,N). An exception to this pattern of behavioral enhancement in “ON” simulations involves visual stimuli placed near the peak of the initial bump in the remember-first condition (Fig. 5B). Under such a scenario, the model predicts that it is more likely that the broader bumps of activity corresponding to the visual stimuli will “merge”, resulting in more errors at the end of the delay period (Fig. 5I). Indeed, as noted above, in the remember-first condition, NB stimulation improved monkey performance in the conditions involving distant distractors (Fig. 5O), however NB stimulation markedly *decreased* performance for visual stimuli appearing at adjacent locations, with excess errors directed to the location of the distractor (two-tailed t-test, *t*_*66*_=2.83, p=0.006). Importantly, this pattern of responses was observed only for the remember-first condition. In the remember-second condition, NB stimulation did not degrade monkey performance for a second visual stimulus appearing at a close distance to the initial distractor (45° condition in Fig. 5R), consistent with the network simulations (Fig. 5L). In addition, we tested the network prediction that in the far-distractor conditions broader bumps generate more stable bump attractors and lower behavioral variability by testing the distribution of angular deviation from the mean endpoint of monkey saccades in correct trials with far distractors. As predicted by the network, this distribution was subtly but significantly narrower under NB stimulation than in the control condition, both in the remember-first (Fig. 5P,Q, Kuiper two-sample test, p=0.01) and in the remember-second conditions (Fig. 5S,T, p=0.001).

**Figure 5.**
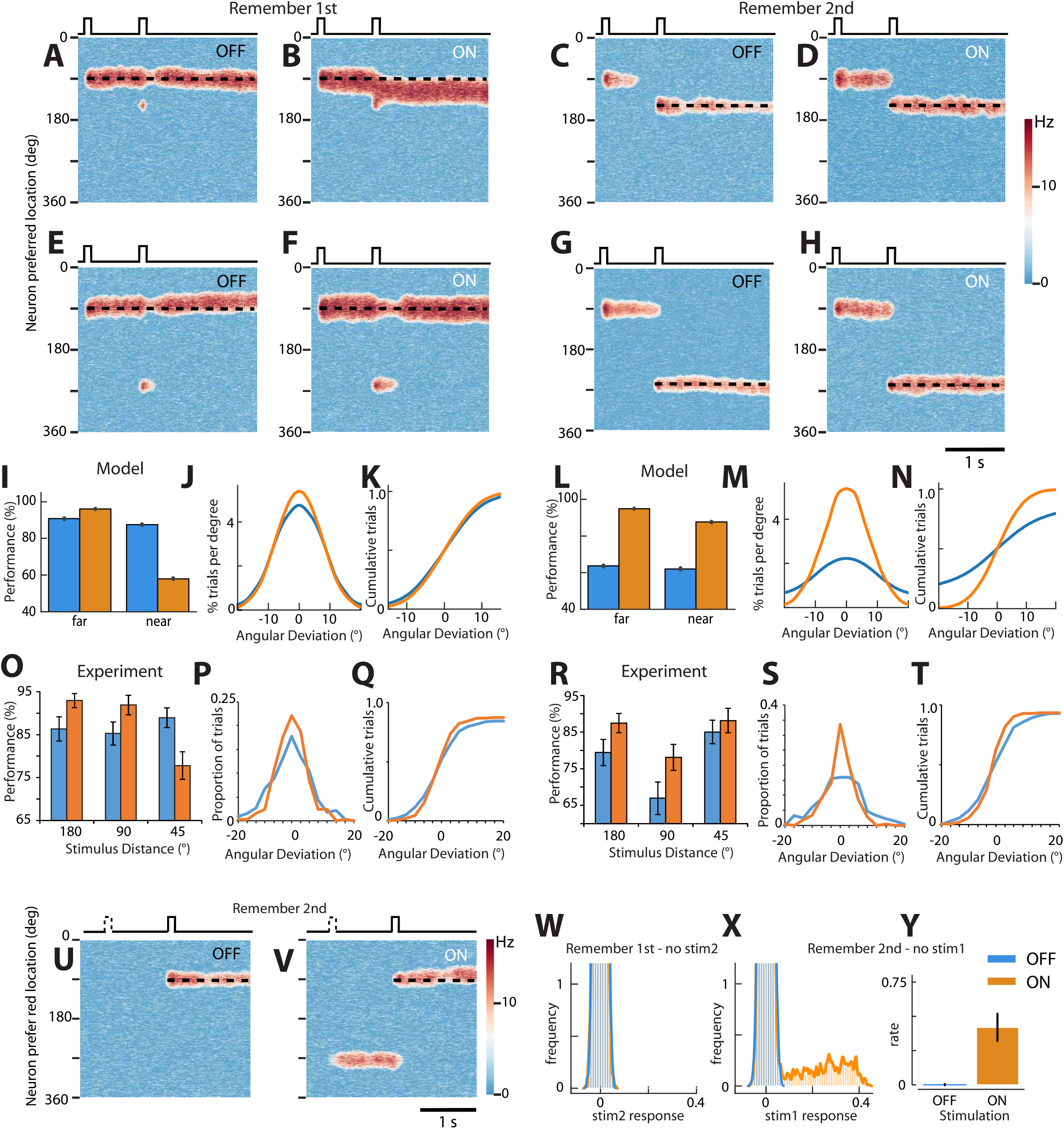
Bump-attractor network behavior. **(A).** Sample simulation showing activity of excitatory neurons in a bump attractor network for the OFF condition in Remember 1st. Abscissa represents time and ordinate represents neurons with preference for different visual stimulus locations, indicated by location varying between 0 and 360°. Activity of neurons with different preference is indicated based on color scale. The first visual stimulus appearance at 90° (indicated by horizontal line on top of the panel) elicits a bump of activity that is maintained during the delay period, after the visual stimulus is no longer present. Appearance of the second visual stimulus at the 150° location does not disrupt the initial bump due to lateral inhibition. The simulated response is the readout of the final location of the bump at the end of the delay period. (**B)**, The ON condition results in a wider bump of activity. As a consequence, when the second visual stimulus appears in a nearby location, it is not inhibited by lateral inhibition and the bumps are more likely to merge, compromising behavior. (**C)** In the remember-second condition, the initial bump of activity terminates and a new bump is maintained after the appearance of the second visual stimulus. (**D)** The ON condition again results in a wider bump of activity, which is more resistant to noise fluctuations, improving behavior. (**E)** For the far conditions, the second stimulus is located at 260°, away from any possible interference with the initial stimulus. (**F)** The ON condition results in a wider bump of activity, which is more resistant to noise fluctuations, improving behavior. (**G-H)** For the remember second condition, the same improvement occurs with NB stimulation (ON condition). (**I)** Remember-first simulations predict a specific pattern of performance as a function of the distance of the distractor to the target, with an increase in performance for the far condition and an impairment in the close condition for ON. Results of 20,000 simulated trials are shown for each condition with a 15° threshold for correct trials. (**J)** Predicted distribution of error in the far condition due to enhanced noise resistance of wider bumps in the ON condition. (**K)** Cumulative distribution of the same data. (**L-N)** Behavioral predictions for remember-second simulations show a general increase in network performance for the ON condition, and narrower distribution of errors in the ON condition. (**O)** Mean monkey performance (and sem) in the remember-first task, for trials grouped by distance between the first and second visual as in Fig. 2I. (**P)** Empirical distribution of angular deviations from mean saccadic endpoint for the far condition, in the remember-first task. (**Q)** Cumulative distribution of the same data. (**R)** Mean monkey performance (and sem) in the remember-second task, for trials grouped by distance. (**S)** Empirical distribution of angular deviations from mean saccadic endpoint for the far condition, in the remember-second task. **(T)** Cumulative distribution of the same data. (**U)** Simulation of the remember-second task with absent first stimulus, under control conditions. (**V)** For the NB-stimulation ON condition, the same simulation generates “phantom” bumps. (**W)** Quantification of phantom bumps in 200 simulations in each condition (ON/OFF). For each neuron and trial, the mean firing rate in the period following the absent second stimulus relative to baseline is divided against the mean response to a preferred stimulus (phantom/evoked activity), and histograms over all neurons and trials (n=102,400) are taken for the ON and OFF conditions. In the remember-first condition, ON and OFF histograms overlap, indicating that phantom bumps following an absent second stimulus are not observed. (**X)** As in W, but for an absent first stimulus in the remember-second task. Phantom bumps are revealed by a slight elevation of the histogram at values >0.1 for the ON condition. (**Y**). Average model unit response following the absent second stimulus, in the ON and OFF conditions.

NB stimulation in the remember-second task yielded another unexpected experimental finding: responses in anticipation of a visual stimulus, even when no visual stimulus was presented at all (Fig. 4F-G). Our behavioral task involved a fixed duration of the fixation interval that the monkey could time. A visual stimulus was most often presented after this interval, however in 20% of the trials no visual stimulus was presented, and the trial continued with the presentation of the “second” visual stimulus at its expected time (see insets in Fig. 4F-G). NB stimulation elicited elevated firing rate in the time interval that the first visual stimulus would have been expected in no-visual stimulus trials (2-way ANOVA with factors tasks and stimulation: main effect of stimulation *F*_*1,53*_=26.9, p=3.4×10^−6^). Such an anticipatory signal was absent without NB stimulation, although presumably the monkey was anticipating a visual stimulus in these trials, too (Fig. 4F-G and S3K). We simulated this anticipation with a non-selective timing signal added to the model at the time of the probable stimulus appearance, as we have documented recently (Barbosa et al., 2020). Phantom bumps of activity were then observed in the model, in the absence of a real visual stimulus. NB stimulation modeled as increased neural excitability elicits a broader, more stable bump but also a more unstable baseline condition, which can quickly develop into a bump when triggered by internal expectation signals (Fig. 5U,V,X,Y). Such phantom bumps were not generated by the model, however, in the remember-first condition, when a stimulus was already actively held in memory and a second stimulus was omitted (Fig. 5W). In the “ON” simulations the network is even more resistant to subsequent stimuli than the control network, and indeed phantom bumps were not observed in neural activity in the remember-first condition (e.g. Fig. S2E).

Network simulations allowed us to address an additional question, regarding the site of action of NB stimulation. In principle, the neural effects we observed might have been entirely the result of changes in upstream, sensory cortical areas that were not active during the delay period of the task, but which were propagated and maintained in the prefrontal cortex. Simulations demonstrated that if that were the case, then a broadening of the bump would only be expected in the cue presentation period, but not in the delay period (Fig. S7), because attractor dynamics impose a fixed bump width in the absence of selective input during the delay period. This outcome was contrary to the experimental results (Fig. 4J) and suggested that NB stimulation must have directly affected working memory circuits, PFC and possibly other areas of the association cortex exhibiting persistent activity, such as the PPC.

### NB Stimulation effects beyond increased firing rate

Neurons that responded to visual stimuli but for which stimulation produced decreased activity (n=16; 7 and 9 from monkeys GR and HE, respectively) were characterized by suppressed firing rate for both the remember-first (Fig. 6A, C) and remember-second tasks (Fig. 6B-D). Firing rate was reduced not only in the fixation period, but also in the visual stimulus presentation period and the delay period that followed it (Fig. 6A, C). Firing rate remained at low levels when the visual stimuli to be remembered were presented out of the receptive field (Fig. 6B, D). NB stimulation flattened the tuning curve of these neurons, too (Fig. 6E). In the group of neurons that did not respond significantly to visual stimuli at all, a general increase in firing rate was observed during NB stimulation, similar to the effects of stimulation on task-responsive neurons (Fig. S4D-E).

**Fig. 6.**
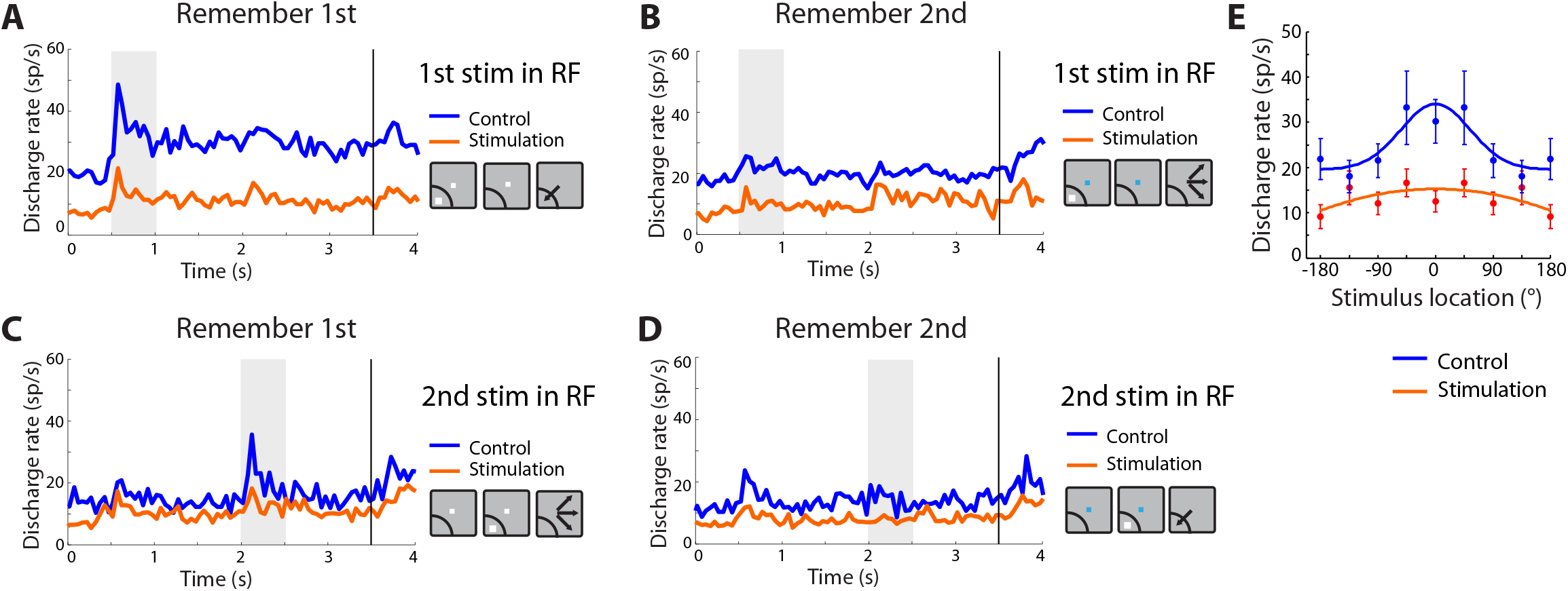
Responses of neurons suppressed by stimulation. Population PSTH represents mean firing rate of neurons whose firing rate during the fixation period decreased during NB stimulation relative to control (n=16 neurons). (**A-B)**. Responses when the first visual stimulus appeared in the receptive field. (**C-D)**. Responses when the first stimulus appeared out of the receptive field. (**E)**. Population tuning curve, obtaining by averaging responses of individual neurons to visual stimuli relative to each neuron’s preferred reference location (depicted at 0°).

Although we emphasize firing rate differences that could account for the behavioral improvements in performance we observed under NB stimulation, alternative mechanisms of working memory have also been proposed, including identifying power in the gamma band of LFP as the critical neural correlate of working memory maintenance (Lundqvist et al., 2018). We therefore examined the LFPs recorded from the prefrontal cortex, in the alpha (8-14), beta (20-35), and gamma (45-100) frequency bands, as these bands were defined in previous studies supporting the latter theory (Lundqvist et al., 2016), and tested if improved working memory performance during the NB stimulation period was the result of increased gamma power. Average spectral power from electrodes in which neurons were also recorded, after subtracting the mean power in the baseline fixation period, is shown in Fig. 7A-B. LFP recordings in the NB stimulation period were characterized primarily by an increase in power in the beta frequency range (Fig. 7C, D). Averaged over the entire delay period and compared across electrodes, the difference reached statistical significance (two-tailed paired t-test, *t*_*113*_=2.77, p=0.0066). Beta-power in the delay period was stable across successive trials in the stimulation block, as was in the inter-trial interval (Fig. 7E-F). No appreciable change in gamma frequency band was evident during stimulation; a decrease in the alpha range was also observed (Fig. 7C-D).

**Fig. 7.**
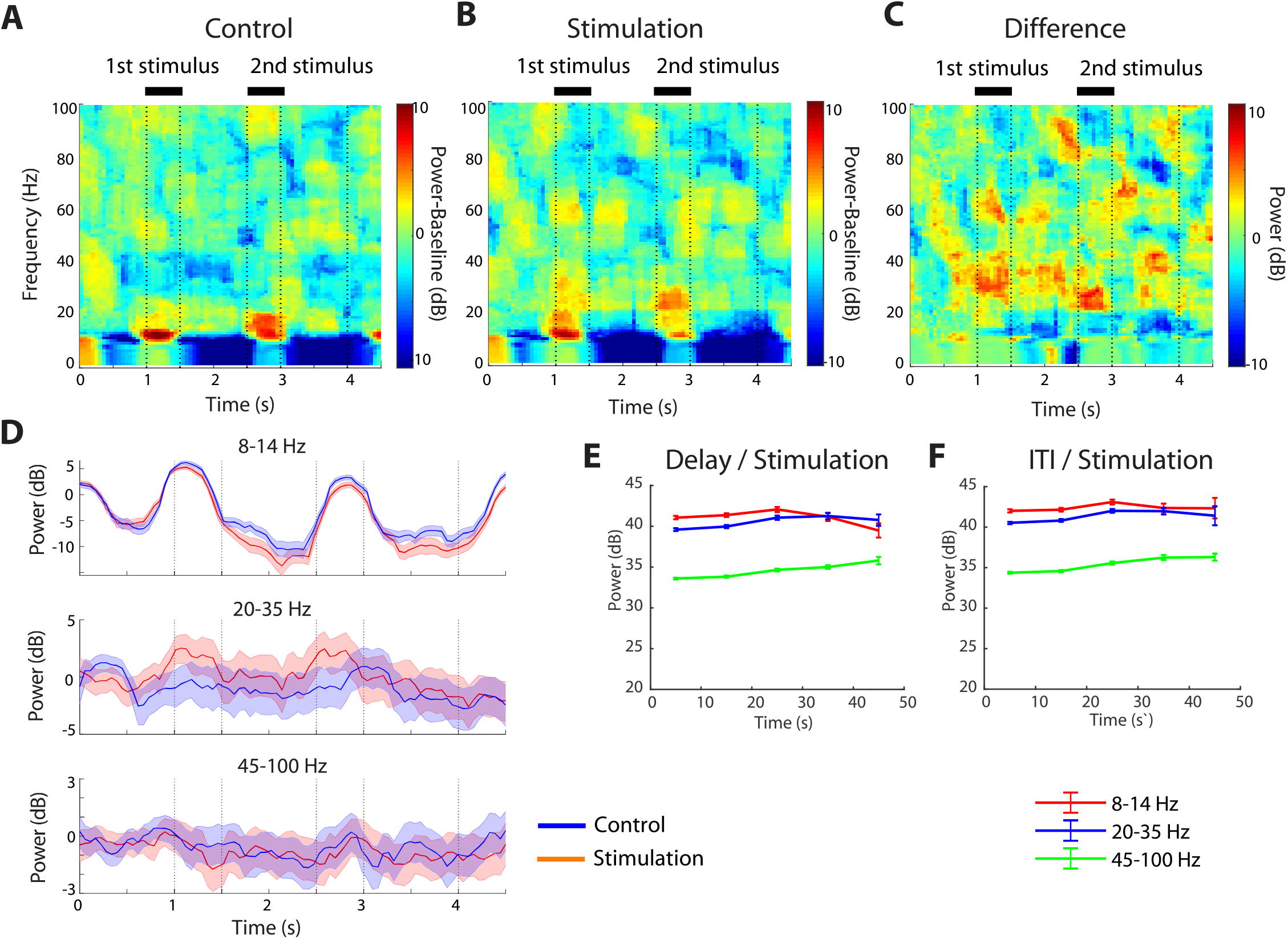
Power Spectrum of Local Field Potentials. **(A)**. Spectral power of Local Field Potentials (LFP) recorded from the prefrontal cortex during the control condition, after subtracting the mean power of the baseline fixation period at each frequency band (n=6096 trials for the control and 7724 for the stimulation condition). Horizontal lines indicate time of stimulus presentations. **(B).** Mean power after subtracting the baseline during the stimulation condition. **(C)**. Difference in power between stimulation and control. Positive values indicate higher power in stimulation condition. **(D)**. Time course of spectral power, after subtracting the baseline, in the alpha, beta, and gamma frequency bands for the control and stimulation conditions. Shaded regions around each line represent 95% confidence intervals, estimated with a bootstrap method. **(E)**. Spectral power in the delay period of the task, in sequential trials of NB-stimulation blocks. Abscissa represents time after the offset of stimulation (as in Fig. 4H). **(F)**. Spectral power in the inter-trial interval.

## Discussion

Our study demonstrated that intermittent NB stimulation improves performance of a working memory task that requires selective maintenance of visual stimuli in memory. Our result are in agreement with recent studies that showed improvements in other cognitive tasks (Blake et al., 2017; Liu et al., 2018, 2017). Taken together, we have now shown working memory improvement by NB stimulation in two different cohorts of monkeys, implanted and tested in two different laboratories (Medical College of Georgia and Wake Forest School of Medicine). We show that improvement in performance was associated with changes in the firing rate of neurons in the dorsolateral prefrontal cortex, most often increasing it. This result was partly consistent with the effects of cholinergic agonists, iontophoresis of which increases activity of prefrontal neurons (Dasilva et al., 2019; Sun et al., 2017; Yang et al., 2013). Conversely, systemic administration of the muscarinic antagonist scopolamine (Zhou et al., 2011) and micro-iontophoresis of muscarinic and nicotinic-α7 inhibitors reduces prefrontal activity (Galvin et al., 2020; Major et al., 2015; Yang et al., 2013). In contrast to these reported drug effects, however, NB stimulation produced an increase that was specific only for some task intervals, notably not affecting the neuron’s preferred visual stimulus response, but increasing firing rate in the fixation interval and the delay intervals over which a visual stimulus was maintained in memory, and amplifying anticipatory responses, prior to the appearance of the visual stimulus. NB stimulation brought about a second unexpected effect, decrease in visual stimulus selectivity. The apparent size of neuronal receptive fields expanded, and spatial selectivity decreased during stimulation.

### Neural tuning implications

The pharmacological studies reviewed above generally suggested that the mechanism through which cholinergic action improves cognitive performance was sharpening of receptive fields, similarly to the effects of dopamine agonists, which at low doses “sculpt” neuronal activity to improve spatial selectivity (Vijayraghavan et al., 2007). Narrower tuning may result in more efficient coding, as the Fisher information per action potential and energy required from the population improves (Fitzpatrick et al., 1997). Broader tuning was only observed in pharmacological studies by cholinergic overstimulation with high doses of carbachol or M1R allosteric inhibitors administered iontophoretically, which reduce prefrontal selectivity in the context of working memory tasks (Galvin et al., 2020; Major et al., 2018; Vijayraghavan et al., 2018). The effect was assumed to correspond to the descending section of an inverted-U function, representing a regime over which cholinergic stimulation will impair performance. It is unknown, however what effective concentration cholinergic stimulation delivers in the prefrontal cortex. Additionally, effects of GABAergic neurons with ascending projections that are concurrently active with cholinergic neurons (Kim et al., 2015; Walker et al., 1989) could not be addressed in iontophoretic studies, and these may also play a role in determining visual stimulus selectivity during NB stimulation.

Broader tuning of individual neurons may not necessarily lead to less efficient coding of visual stimulus information (Ma et al., 2006; Zhang and Sejnowski, 1999). One existing class of models enables exploration of the impact of broadened tuning on working memory accuracy (Compte et al., 2000; Wimmer et al., 2014). In this model, called the bump attractor, neurons are connected to each other depending on visual stimulus selectivity. A visual stimulus evokes a response in neurons with similar selectivity, which in the model resembles a bump. The network behaves as a continuous attractor (Seung et al., 2000), allowing activity to persist even when the original visual stimulus is no longer present but making the storage of parametric quantities sensitive to internal noise fluctuations. The activation of a larger population of neurons by a single visual stimulus, consistent with decreased neuronal selectivity, results in a broader bump of activity rendering the network more resistant to noise fluctuations that perturb memory maintenance. At the same time, performance was predicted to be compromised in conditions involving distracting visual stimuli appearing in nearby locations. These predictions were born out by the behavioral results (Fig. 5). We also observed prefrontal responses in anticipation of visual stimuli that did not appear, which is consistent with more excitable attractor networks, in which spurious activation during spontaneous activity may create “phantom” bumps. We should also note, however, that improvement by NB stimulation was not uniform across conditions. NB stimulation degraded performance when ipsilateral stimuli were held in memory and when distractors were present. The broadening of neuronal tuning is not without costs, therefore, and the pharmacological studies mentioned above likely reflect some of these decreases in performance.

Although we emphasize the similarity between the modeling and neural results, it is also important to point out the limitations of the model. The model relies on a change in the strength of inhibitory and excitatory connections to simulate the task in which the monkey remembered the first or second stimulus. How this is performed in the brain is currently unknown, and local-circuit neuromodulation is only one of several possible mechanisms. One possibility is that sensory inputs of unwanted distractors get suppressed (Nakajima et al., 2019), so they do not even reach memory storage centers. Alternatively, there could be changes in storage circuits, such that incoming inputs are less likely to enter the memory state if there is already one memory stored. This may happen through large-scale brain-area interactions (Sakai et al., 2002) or by virtue of the neuromodulation of local-circuit synaptic interactions, as demonstrated in attractor networks of working memory (Compte et al., 2000; Durstewitz et al., 2000).

### Areal specialization

We should note that the effects of cholinergic agents in sensory areas are markedly different from those in the prefrontal cortex. Agonist administration in the primary visual cortex specifically enhances responses during visual stimulus presentation, and attended over unattended ones (Herrero et al., 2008). Similarly, optogenetic phasic stimulation of cholinergic neurons has been shown to improve visual perceptual discrimination by increasing firing rate during the period of visual stimulus presentations (Pinto et al., 2013). Neuronal responses within the Nucleus Basalis often signal novelty or surprise (Zhang et al., 2019) and widely distributed projections of these neurons appear to play different roles at different parts of the cortex. Stimulation results also argued directly against the possibility, of the effects of stimulation operating exclusively at the level of the sensory cortex (Fig. S7). On the other hand, NB stimulation is likely to activate other areas of the association cortex, in addition to the prefrontal cortex, including the posterior parietal association cortex (Mesulam et al., 1983). PPC neurons exhibit persistent activity in similar tasks, and its activation by NB stimulation is both consistent with our experiment results, and their known role in cognitive performance (Riley and Constantinidis, 2016).

### Potential mechanisms of action

It is important to point out that our protocol of NB stimulation can improve cognitive performance in healthy adult monkeys, but only when administered in an intermittent fashion. Optimal stimulation parameters involve stimulation for 15-20 seconds per minute, and at a rate that delivers approximately a total of 1200 pulses of stimulation per minute (Liu et al., 2017). The result suggests that continuous stimulation may lead to depletion of acetylcholine reserves, a phenomenon consistent with the known impairment of working memory caused by acetylcholine depletion in the prefrontal cortex (Croxson et al., 2011). Prolonged systemic administration of cholinergic agents may have a similar effect, which would account for its loss of effectiveness in Alzheimer’s patients.

The performance improvement induced by NB stimulation depends on acetylcholine release, as cholinergic inhibitors abolish the performance benefits of intermittent stimulation (Liu et al., 2018). The effect on different cholinergic receptor subtypes is likely complex, however. Nicotinic α7 agonists produce predominant excitatory effects (Yang et al., 2013) whereas suppressed firing can be induced with high doses of muscarinic m1 receptor agonists (Vijayraghavan et al., 2018). In rodent PFC, m1 receptors excite GABAergic interneurons (Tikhonova et al., 2018), suggesting that NB stimulation triggers complex network effects.

In addition to cholinergic efferents, GABAergic ascending projections are likely activated by our protocol of NB stimulation and play a more physiologically relevant role in the modulation of responses in target areas (Walker et al., 1989; Zaborszky et al., 2018). In rodents, GABAergic projections from the basal forebrain are electrically coupled and responsible for precisely-timed, rhythmic entrainment of cortical neurons, which is evident in increase of gamma-band oscillations when they are selectively activated (Kim et al., 2015; McKenna et al., 2013). We do note, however, that the aggregate effect of our NB stimulation did not increase LFP gamma-band power in the prefrontal cortex, in agreement with recent NB stimulation results in the rodent auditory cortex (Azimi et al., 2020). The combination of judicious pattern of stimulation and recruitment of the full apparatus of ascending modulatory control appears critical for the success of the intervention.

## Acknowledgments

Research reported in this paper was supported by the National Institute of Mental Health of the National Institutes of Health under award numbers R01 MH097695 and RF1 AG060754. DB and AC were supported by the Spanish Ministry of Science and Innovation and European Regional Development Fund (Ref: RTI2018-094190-B-I00) and by the CERCA Programme/Generalitat de Catalunya. We wish to thank James Daunais, Kathini Palaninathan, Kristopher Bunting, George Pate, Austin Lodish, Sihai Li, and Junda Zhu for technical help; Alvin Terry for guidance with the design of the study; John Murray and Joost Maier for valuable comments on the manuscript.

## Author contributions

C.C. and D.T.B. designed the experiments. R. L., C.C., X.-L.Q., and D.T.B. performed surgeries. X.-L.Q. and C.C. conducted behavioral and neurophysiological experiments. X.-L.Q., B.S., and C.C. performed data analysis. A.I.V. conducted immunohistochemistry experiments. D.B. and A.C. performed model simulations. C.C., X.-L.Q. and D.T.B. wrote the manuscript, with input from all authors.

## Methods

Two male, rhesus monkeys (*Macaca mulatta*) weighing 7-10 kg were used in this study. All experimental procedures followed guidelines by the U.S. Public Health Service Policy on Humane Care and Use of Laboratory Animals and the National Research Council’s Guide for the Care and Use of Laboratory Animals and were reviewed and approved by the Wake Forest University Institutional Animal Care and Use Committee.

### Surgery and neurophysiology

A 20-mm-diameter recording cylinder was implanted over the dlPFC (Fig. 1). A second cylinder was also implanted over the PPC of each monkey at the same time, but this was not used in the current experiment. Extracellular activity of single units was recorded from areas 8a and 46 of dlPFC. The anatomic location of electrode penetrations was determined on the basis of MR imaging. Recordings were obtained with arrays of two to four microelectrodes in the cylinder. These were epoxylite-coated tungsten electrodes with a 250 μm diameter and 1-4 MΩ impedance at 1 kHz (FHC, Bowdoin, ME). The electrical signal from each electrode was amplified, band-pass filtered between 500 Hz and 8 kHz, and recorded with a modular data acquisition system at 25-μs resolution (APM system; FHC, Bowdoin, ME). Waveforms that exceeded a user-defined threshold were sampled at 25 μs resolution, digitized, and stored for off-line analysis. LFP recordings were obtained from the same electrodes by splitting the signal, filtering between 0.5 Hz and 100 Hz, and acquiring data at a 500 Hz sampling rate.

### Deep Brain Electrode Implantation and Stimulation

Once the head-cap and recording cylinders had been implanted, a second surgery was performed to implant the stimulating electrode. Based on MR imaging, stereotaxic coordinates were obtained for targeted implantation. The animals were implanted unilaterally (one in the left, and one in the right hemisphere) at 8 mm lateral, 16 mm anterior interaural, and 29 mm below the cortical surface in a vertical penetration. The lateral and anterior coordinates, and depth, were chosen to correspond to the center of the anterior portion of the Nucleus Basalis of Meynert, which would contain the highest density of projections to the prefrontal cortex (Gielow and Zaborszky, 2017; Mesulam et al., 1983). A small cylindrical titanium chamber (5-mm inner diameter and 7-mm outer) was mounted on the cranium and chamber was encased in bone cement, in continuity with the existing head-cap. A 26 ga. sharp hypodermic guide tube was lowered and the tip advanced 5 mm below the dura mater. The stimulation electrode was inserted into the guide tube, and a stylus was used to push it to the appropriate depth. The guide tube was then raised while the stylus depth maintained. The chamber was evacuated of fluid, flushed with ceftriaxone, and thereafter fluid evacuated a second time. Silicone was poured into the chamber to seal the fenestrations in the skull and the inside of the chamber. The rear end of the electrode could be continuously visualized to confirm proper depth. The electrode was fixed in depth with a drop of cyanoacrylate. One week after the surgery, the animals returned to behavioral studies. Placement of the electrode was verified with CT scanning, after implantation, in one animal.

The stimulation pulses were created by an isolated pulse stimulator (Model 2100, A-M Systems, Sequim WA), which was controlled by custom programed software, written on the MATLAB platform. Impedances of electrodes were checked monthly during experiments. Intermittent stimulation was applied for 15 seconds at 80 pulses per second, followed by approximately 45 seconds with no stimulation. Stimulation was applied in the inter-trial interval, after a trial had completed, and a new trial began after stimulation had elapsed.

Stimulation electrodes were custom manufactured in our laboratory based on published specifications (McCairn and Turner, 2009). Conductors were 50 μm Pt/Ir, Teflon-insulated wire (A-M systems, Seattle, WA) embedded within a 30 ga. hypodermic tube, which was encased in a 28 ga. polyimide sheath. The wire extended from the end of the sheath into the brain tissue by roughly 1 mm, and the last 0.7 mm of insulation was stripped to achieve impedances of 5-10 kOhm at 1 kHz. The far end of the electrode was soldered to an extracranial connector fixed on the chamber outer wall. Prior experiments on electrode placement tested the effects of short periods of stimulation on EEG desynchronization (Liu et al., 2017). Stimulation was delivered with biphasic, negative first, unipolar 200 μA pulses with 100 μs per phase, and 80 Hz pulses were delivered for 100 ms. This resulted in LFP desynchronization obtained through the stimulation electrode when the electrode was at a depth corresponding to the atlas position of Nucleus Basalis. In prior experiments, an electrode movement vertically in either direction of more than 1 mm was adequate to make desynchronization not possible using the same protocol (Liu et al., 2017). LFP recordings obtained through the stimulating electrode used the same filtering and sampling parameters as the recording electrodes (band pass filtering between 0.5 – 100 Hz, sampling at 500 Hz).

### Behavioral tasks

The monkeys faced a computer monitor 60 cm away in a dark room with their head fixed. Eye position was sampled at 240 Hz, digitized, and recorded with an infrared eye position tracking system (model RK-716; ISCAN, Burlington, MA). The visual stimulus presentation and behavior monitoring were controlled by in-house software (Meyer and Constantinidis, 2005) implemented in the MATLAB computational environment (Mathworks, Natick, MA).

The tasks used in the present study were variations of the Oculomotor Delayed Response task, but involving two visual stimuli appearing in sequence, requiring the monkey to remember and make an eye movement to the location of either the first or the second visual stimulus (Fig. 2). The monkeys were trained to saccade to the location of the remembered visual stimulus according to the color of fixation point. After the animals fixated at a white/blue square (0.2° in size) located at the center of the monitor for 1 second, two white squares (1.5° in size, 125 cd/m^2^ in luminance, 99% Michelson contrast) were displayed sequentially for 0.5 s, with a 1 s intervening delay period. The first visual stimulus was displayed at one of eight possible locations arranged along a circular ring of 12° eccentricity, with a 45° angular separation between neighboring visual stimuli. The monkeys were trained with stimulus appearances at every possible location, however, during recordings the first stimulus appeared at one of these eight locations and its diametric location. This was followed by a second visual stimulus, which was displaced 0, 45, 90, or 180° relative to the first. Two additional, “null” conditions were included, in which either the first or second stimulus presentation was omitted, so that there were 10 trial types in total, and these were used with equal frequency. After a second delay period of 1s, the monkeys were required to saccade to the location of the first visual stimulus if the fixation point was white in color (remember-first condition), and to the location of the second visual stimulus if the fixation point was blue (remember-second condition). The monkeys were rewarded with juice after making a correct saccade. Deviating gaze beyond a 3°-radius fixation window led to the immediate termination of the trial without reward.

At the beginning of each recording session, we first ran the ODR task with a single stimulus, as a way to approximately map the receptive fields of neurons isolated from our electrodes. Then the remember-first and remember-second task began. To minimize the uncertainty about the visual stimulus to be remembered, the remember-first and remember-second conditions were presented in blocks of trials. The animal was required to perform ten correct trials of the remember-first task, before the task alternated to the remember-second condition. During sessions of NB stimulation that were delivered in block mode (trials with and without stimulation collected during the same daily session), approximately 60 trials were collected for each block, which therefore involved 6 alterations between the remember-first and remember-second rule.

### Behavioral Performance

We calculated behavioral performance as the percentage of trials that resulted in correct saccades into the target window, a 7° circle around the center of the visual stimulus. Trials that were aborted prior to end of the second delay period (due to premature saccades, or blinks) were not included in this analysis. Performance in a single daily session was used as a unit of analysis in Figures 2 and 5. A daily session included approximately 120 trials.

### Neural Data Analysis

All data analysis was implemented with the MATLAB computational environment (R2012-2019, Mathworks, Natick, MA). Recorded spike waveforms were sorted into separate units using an automated cluster analysis relying on the KlustaKwik algorithm (Harris et al., 2000), which relied on principal component analysis of the waveforms. Mean firing rate was then determined in each task interval. Neurons responsive to the visual stimuli were identified, evidenced by a significant increase in firing rate in either the cue period or delay period relative to the baseline fixation period (paired-t test, evaluated at the 0.05 significance level). These neurons were used for most analyses in the main text, however analyses including all neurons were also performed. Neural data from correct trials were used only in these analyses. Units were identified as broad-action potential, Regular Spiking (RS) neurons or narrow-action potential, Fast Spiking (FS) neurons, based on the width of the action potential. This was defined as the duration between two positive phases of the waveform, using a criterion value of 0.55 ms, as described before (Qi et al., 2011).

We identified neurons with a significant excitatory or inhibitory effect of stimulation by comparing baseline firing rate during the fixation period between the control and stimulation conditions (evaluated with a t-test at the p<0.05 level). To study separately effects of stimulation on the remember-first and remember-second task, we compared firing rates across conditions by performing a 2-way ANOVA with factors tasks and stimulation. We repeated this 2-way ANOVA for other task intervals, including the visual stimulus presentation and delay period. In a similar fashion, a 3-way ANOVA was performed in order to determine the main effect of task, location of first visual stimulus, and location of second visual stimulus on firing rate. This analysis was performed in a time-resolved fashion, in successive 250 ms windows spanning the entire trial.

The Fano factor of a neuron’s discharge rate (defined as the variance divided by the mean) was estimated in different task periods. Data for each neuron and condition were treated separately. Spike counts were computed in a 100-ms sliding window moving in 20-ms steps. The method computes the variance and mean of the spike count across trials and performs a regression of the variance to the mean (Qi and Constantinidis, 2012). This slope of this regression represents the Fano Factor reported here.

We quantified selectivity for different visual stimulus location using a Selectivity Index (SI) defined as (Max-Min)/(Max+Min) where Max and Min represent the firing rate corresponding to the best and worst stimulus location for each neuron, determined independently for the cue and delay periods, and calculated over the entire cue/delay period. SI values were also determined independently for the control and NB-stimulation conditions. Discriminability between visual stimulus conditions does not only depend on mean firing rates, on which SI depends, but also their variance. We therefore compared the full distributions of firing rates between the worst and best condition using a Receiver Operating Characteristic analysis. This was performed in a time-resolved fashion, in successive 250 ms windows.

The endpoint of saccades towards the targets were analyzed in correct trials. For each monkey and visual stimulus condition, the mean saccadic endpoint was calculated for the target location (using circular mean statistics). We then computed the angular deviation of the saccadic endpoint in every trial, relative to the mean saccadic endpoint for that condition. We applied the Kuiper two-sample test, a circular analogue of Kolmogorov-Smirnov test, to compare the distributions of angular deviations around the mean, with and without stimulation.

We used the FieldTrip toolbox (Oostenveld et al., 2011) for preprocessing analysis and Chronux package (Bokil et al., 2010) for time-frequency analysis. The Local Field Potential signal obtained through the stimulation electrode was determined at rest and following 6 seconds of intermittent stimulation. Multiple sessions were obtained to determine desynchronization in this fashion. For power analysis of LFP signals from the recording electrodes, a band-pass filter (0.5-200 Hz) was used. We removed line power (60 Hz) and other artifacts from each electrode and trial in the LFP signal, if present. We then used a multi-taper method to perform a power spectrum analysis of LFP. Power spectra were constructed, and plotted after subtracting the mean power of the baseline fixation period at each frequency. We then compared the LFP power at each frequency between the control and simulation conditions. We also analyzed the LFP power at different frequency bands defined as alpha (8-14 Hz), beta (20-35 Hz) and gamma (45-100 Hz) after subtracting the mean power. We obtained estimates of the variability of these measures by using a bootstrap technique of randomly selecting trials with replacement, and repeating the power spectrum estimation 1,000 times.

### Bump-attractor model

We simulated a bump attractor network model in a firing-rate neuron formulation (Edin et al., 2009; Wimmer et al., 2014). The network consists of fully interconnected excitatory (N_E_=512) and inhibitory neurons (N_I_=512). Neurons present selectivity for specific angles (*θi* for i=1..N) on a ring of fixed eccentricity in the visual scene, and neurons with similar selectivity are more strongly connected than neurons coding for distant locations. This is modeled with connectivity strengths *w*_*ij*_ that follow von Mises distributions (*W*_*ij*_ = *Aexp* (*KCOS*(*θ*_*i*_ - *θ*_*j*_)) with *K*_*E*_ = 45 and *κ*_*I*_ = 0.3 for excitatory and inhibitory connections, 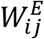 and 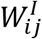, respectively, and a normalization factor *A* set so that 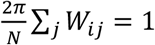. These normalized connectivities are then scaled by a connectivity strength parameter *G*. Visual stimuli are modeled as an external current applied for 100ms to excitatory neurons during stimulus presentation, with intensity peaking at the location of the stimulus according to a von Mises distribution (as above) with *κ*_*stim*_ =40 and intensity scaling *G*=9.4. In addition, we simulated an “expectation signal” concomitant with the stimulus presentation as an additional non-specific input to the excitatory network with strength 1.2. This represented an internal signal that predicted the regularly timed stimulation presentation events in the task, and it was relevant to simulate the emergence of phantom bumps when the stimulus was not presented (Fig. 5). Upon the extinction of the visual stimulus input, reverberatory activity maintains the information in the form of self-maintained selective elevated activity, a bump attractor (Fig. 5A-H). The equations that define the evolution of the rates of excitatory 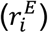 and inhibitory 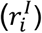 neurons in our model are:

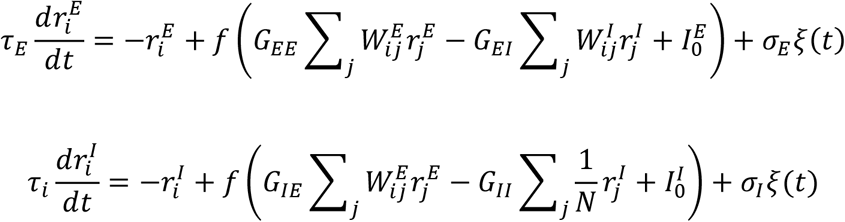

with neuronal time constants *τ*_*E*_=20 ms and *τ*_*I*_= 10 ms; *G*_*EE*_, *G*_*EI*_, *G*_*IE*_, and *G*_*II*_ synaptic strengths (values below); 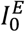 and 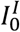 untuned background inputs (values below); and *σ*_*E*_=9.2 and *σ*_*I*_=6.6 amplitudes of uncorrelated Gaussian white noise input *ξ*(*t*). During the stimulus presentation period, both populations receive the extra input of the stimulus current described above. The model transforms currents into rates through a neural transfer function *f*(*I*) = 0 for *I*<0, *f*(*I*)=*I*^*2*^ for *0*<*I*<*1*, and 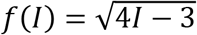.

We modeled the remember-first or remember-second conditions of the experiment with slightly different connectivity parameters and input (similar to Compte et al. 2000) to reflect top-down influences in the specific blocks of these conditions. Specifically, we modulated *G*_*EE*_ (1st=0.068, 2nd= 0.064), *G*_*II*_ (1st=0.13, 2nd=0.01196), *G*_*EI*_ (1st=0.13, 2nd=0.1482), and *G*_*IE*_ (1st=0.042, 2nd= 0.045), and 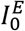 (1st=−3.5, 2nd=−2).

To model the Nucleus Basalis stimulation (ON condition), we assumed that release of ACh in PFC would result in blockade of hyperpolarizing intrinsic currents in excitatory neurons, so we increased the excitability of this population through an increase of 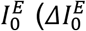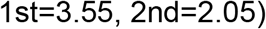 For all conditions 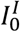 was fixed at 0.5.

### Immunohistochemistry and Fluorescence Imaging

Free floating 50-μm sections of fixed monkey brain were stained with Anti-ChAT antibody (MilliporeSigma, MAB5270), Goat Anti-Mouse-Biotin-SP antibody (Jackson ImmunoResearch Laboratories, 115-067-003), Streptavidin-HRP (PerkinElmer, NEL750), and SuperGlo Green fluorescein tyramide (Fluorescent Solutions, FS101). Stained sections were mounted on slides and z-stack images were collected at 10X and 20X magnification using a Zeiss AxioImager microscope. The coronal section displayed in Fig. 1C was corresponded to a plane approximately 16 mm anterior to the interaural line.

## SUPPLEMENTAL FIGURES

**Fig. S1.**
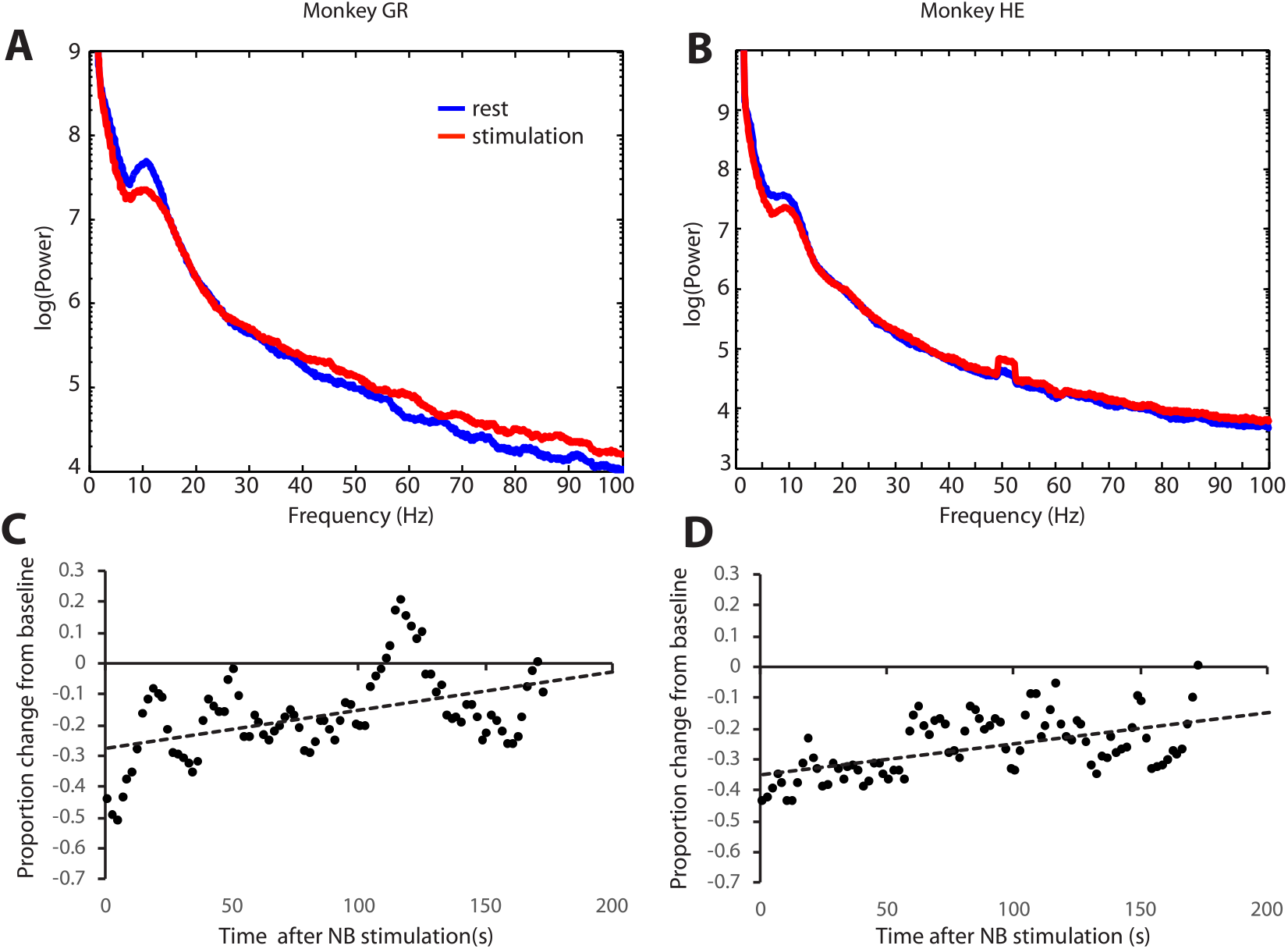
Supplement to Figure 1. Local Field Potential Desynchronization. **A-B.** Power Spectrum of Local Field Potential recorded from the implanted electrode during rest and following 80 Hz stimulation in the two animals. Same data as in Fig. 1A, plotted with an extended X axis, up to 100 Hz. **C-D.** Time course of decrease in power relative to baseline in the 5-15 Hz frequency range. Individual points represents power decrease relative to baseline at successive 6-second intervals. Dotted traces represent regression lines (slope β=0.123 % s^−1^ for subject GR, β=0.104 % s^−1^ for subject HE).

**Fig. S2.**
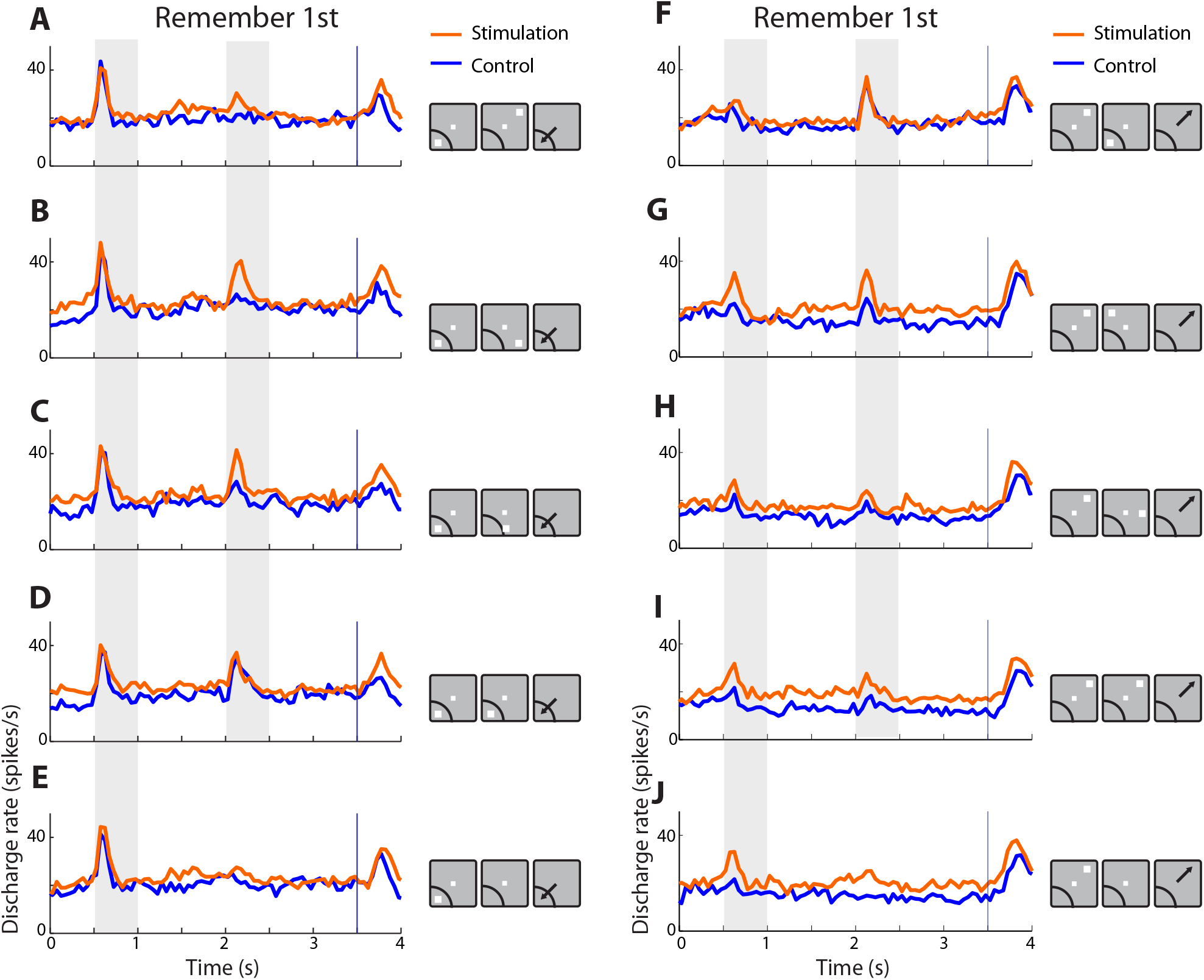
Supplement to Figure 4. Neuronal responses for all stimuli in remember-first task. Mean firing rate of neurons with significant increase in activation by NB stimulation (n=54 neurons). **A-E**, conditions involving presentation of the first stimulus in the receptive field. Gray bars represent times of stimulus presentations. Insets to the right of PSTH represent location of the stimuli relative to each neuron’s receptive field; results from neurons with different receptive field locations have been averaged together, but only one stimulus location is indicated. **F-J**. Mean firing rate in conditions involving presentation of the first stimulus away from the receptive field.

**Fig. S3.**
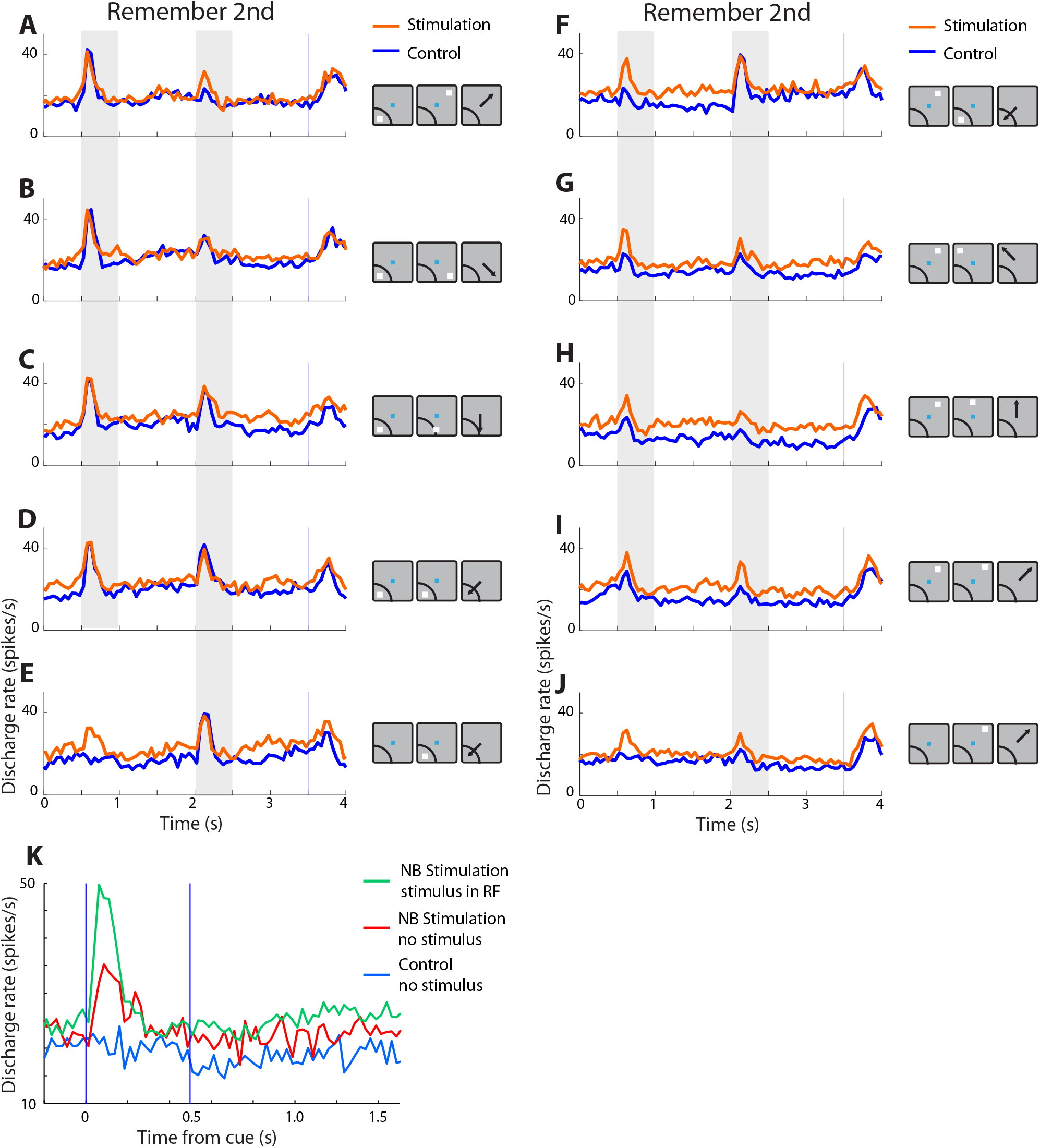
Supplement to Figure 4. Neuronal responses for all stimuli in remember-second task. Mean firing rate of neurons with significant increase in activation by NB stimulation (n=54 neurons). **A-D**, conditions involving presentation of the first stimulus in the receptive field. Gray bars represent times of stimulus presentations. Insets to the right of PSTH represent location of the stimuli relative to each neuron’s receptive field; results from neurons with different receptive field locations have been averaged together, but only one stimulus location is indicated. **E.** Null condition, involving no first stimulus, followed by a second stimulus in the receptive field. **F-I**. Mean firing rate in conditions involving presentation of the first stimulus away from the receptive field. **J**. Null condition, involving no presentation of the first stimulus, followed by a second stimulus away from the receptive field. **K**. Average firing rate from conditions with stimulus in the receptive field is contrasted with the null conditions and conditions with no stimulus in the receptive field in higher resolution.

**Fig. S4.**
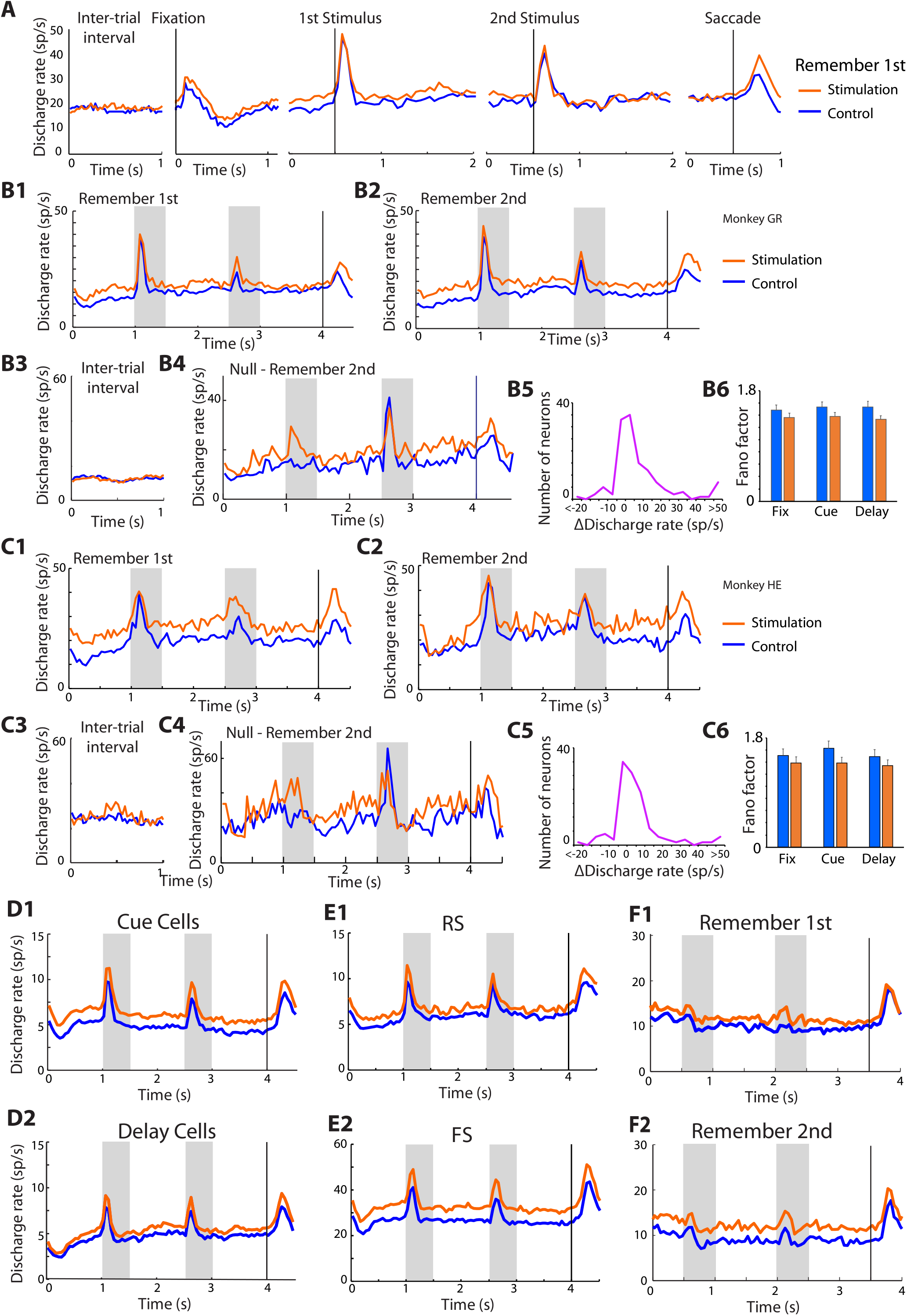
Supplement to Figure 4. Consistent effects of stimulation across tasks, subjects, regular spiking and fast spiking neurons, and neurons without elevated responses to visual stimuli. **A.** Results from the remember-first task are shown, for neurons with significant increase in activation by NB stimulation (n=54 neurons). Conventions are the same as Fig. 4A. Mean firing rate with and without stimulation is shown during the inter-trial interval, fixation interval, first stimulus presentation involving the best stimulus of each neuron, second stimulus presentation involving the best stimulus of each neuron, and saccade towards best stimulus. **B1**. Mean firing rate with and without NB stimulation is shown for the remember-first task for neurons with significant increase in activation by NB stimulation for monkey GR (n=41 neurons). Conditions with the first stimulus appearing in the best location have been pooled together. **B2.** As in B1, for the remember-second task. **B3**. As in B1, for the inter-trial interval. **B4**. As in B1, for the remember-second task, when no stimulus appeared at the first stimulus interval (null condition, as in Fig. 4F). **B5**. Distribution of firing rate changes in fixation period after NB stimulation (as in Fig. 3F) for the same monkey. **B6.** Mean Fano factor values (as in Fig. 3G) for the same monkey. **C1-C6**. As in B1-B6, for monkey HE (n=13 neurons). **D1**. Neurons responsive in the task with cue responses, but no delay period activity (n=22). **D2**. Neurons responsive in the task with delay period activity (n=32). **E1**. Mean firing rate of all Regular Spiking, (putative pyramidal) neurons in the sample (n=154 neurons). **E2**. Mean firing rate of all Fast Spiking (putative interneurons, n=64). **F1.** Mean firing rate of neurons which, in the control condition, did not respond with a significant increase to the appearance of the stimulus, or the delay period following it (n=118 neurons) Responses in the remember-first task are shown. **F2.** Mean firing rate of same neurons as in panel F1, for the remember-second task.

**Fig. S5.**
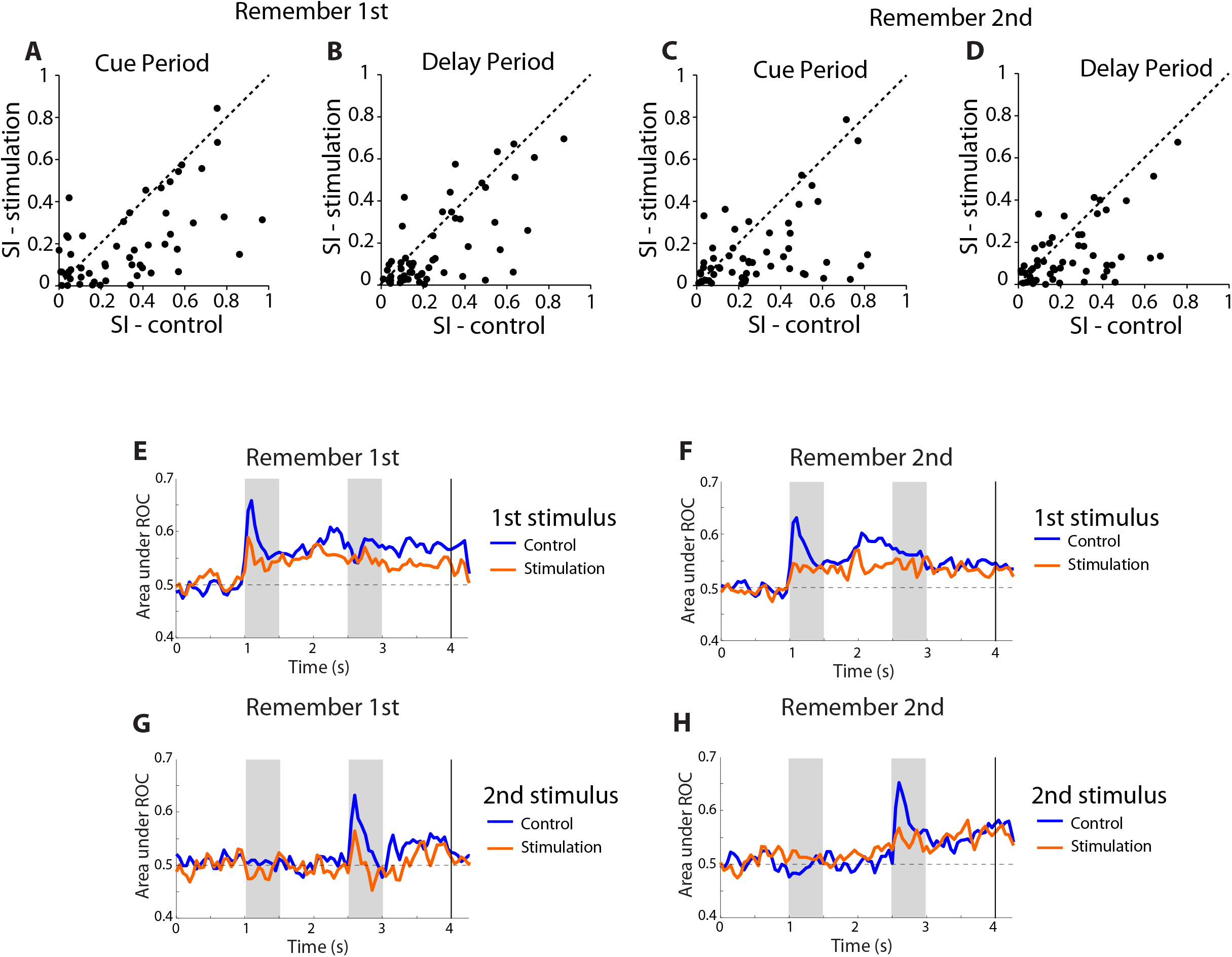
Supplement to Figure 4. Selectivity Index and ROC analysis. **A**. Selectivity Index (SI) for the cue (first stimulus) period of the remember-first task. Each data point represents the SI of a single neuron during the stimulation (ordinate) and control conditions (abscissa). SI is defined as (Max-Min)/(Max+Min) where Max and Min represent the averaged firing rate during the cue period eliciting the maximum and minimum response. Most data points fall below the diagonal (dotted line). **B.** SI values during the delay period of the remember-first task. **C**. SI values during the cue period of the remember-second task. **D**. SI values during the delay period of the remember-second task. **E**. Mean value of area under ROC curve across neurons, in the control and stimulation conditions. Each point represents the ROC value determined based on the distribution of firing rates elicited in trials when the first stimulus appeared in the best location for each neuron, vs. trials in which the first stimulus appeared in the worst location for the neuron. This calculation was performed at different time points across the trial, to produce the two traces. **F**. Area under the ROC curve for the first stimulus in the remember-second task. **G**. Area under the ROC curve for the second stimulus in the remember-first task. **H**. Area under the ROC curve for the second stimulus in the remember-second task.

**Fig. S6.**
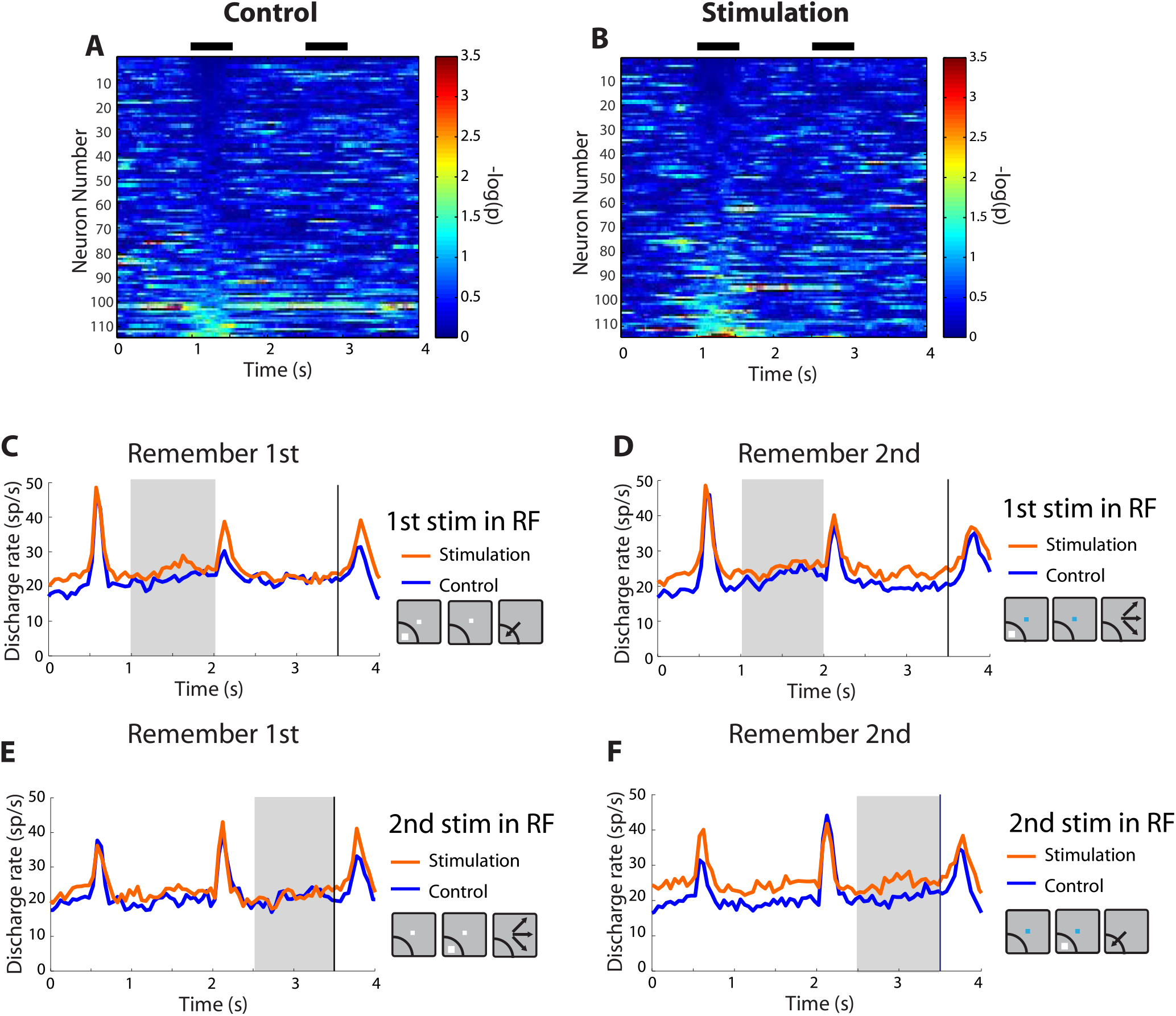
Supplement to Figure 4. Representation of task and stimulus information. **A.** Main effect of task is plotted for each neuron and time point in a 2-way ANOVA with cue stimulus location and task (remember-first / remember-second) as factors, in the control condition. Color represents negative logarithm of p-value of main task effect. **B.** Same as in panel A, under stimulation condition. **C**. Mean firing rate averaged across all stimulus conditions of the remember-first task in which the first stimulus appeared in the receptive field. Gray bar represents delay period activity following this stimulus. **D.** Mean rate for conditions of the remember-second task in which the first stimulus appeared in the receptive field. **E.** Mean firing rate averaged across all stimulus conditions of the remember-first task in which the second stimulus appeared in the receptive field. **F**. Mean firing rate averaged across all stimulus conditions of the remember-second task in which the first stimulus appeared in the receptive field (n=54 neurons).

**Fig. S7.**
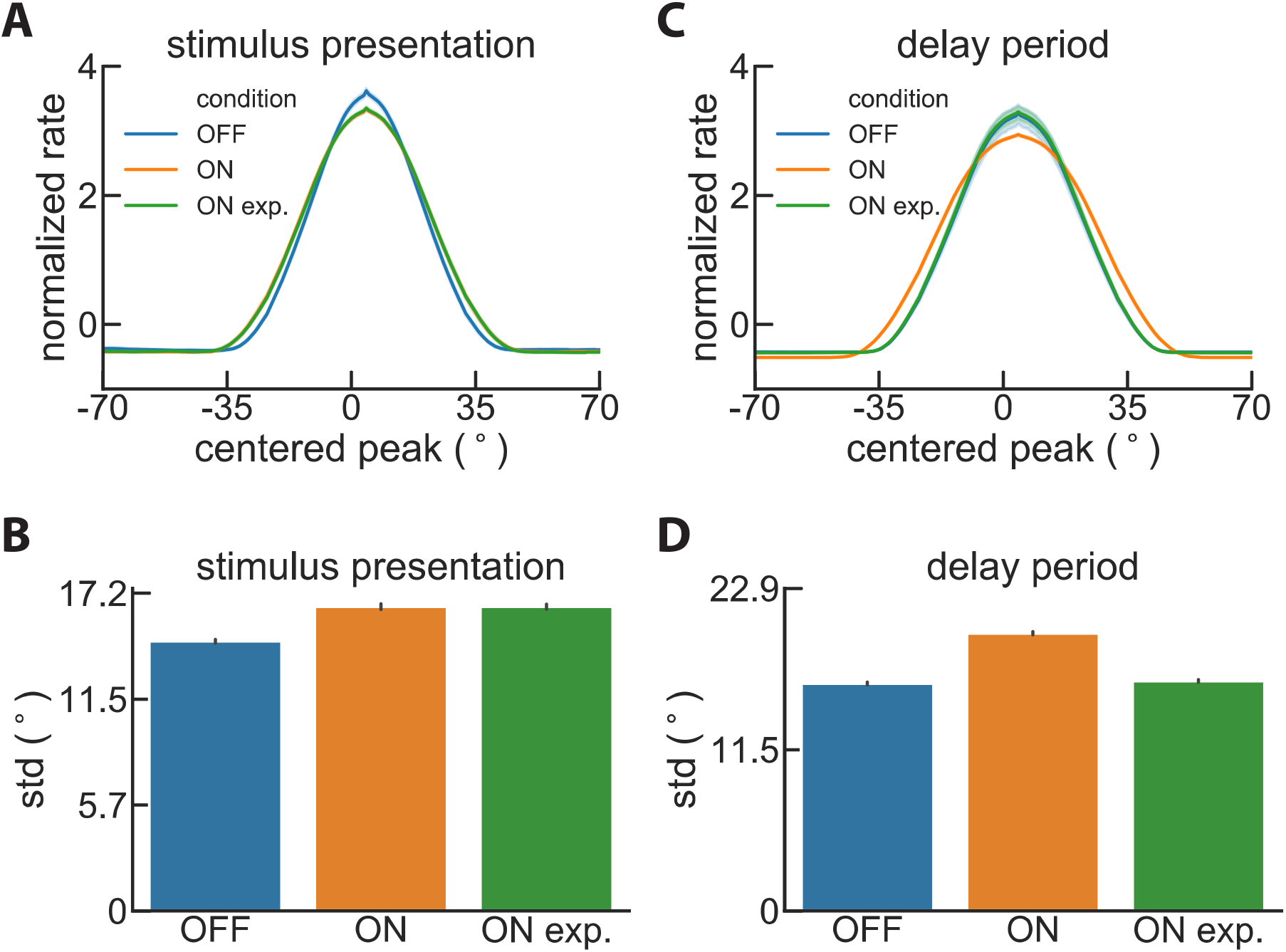
Supplement to Figure 5. Activity bump during NB stimulation. **A.** Results of simulating the effect of NB stimulation as affecting the PFC circuit itself, as a constant increase in the excitability of the working memory circuit (ON), and as affecting primarily the areas upstream from the working memory circuit, so all that this circuit sees is an increased expectation signal (ON expect) during stimulus presentations. The width of the bump is plotted during the stimulus presentation. **B.** Standard deviation of the bump shown in panel A. An in increase of bump width during the stimulus presentation is evident for the ON expect condition (as in the data). **C-D**. Simulation results for the delay period. Conventions are the same as in panels A-B. No change in bump width compared to OFF during the delay period is now evident for the ON expect condition, contrary to data. This is because attractor dynamics imposes a fixed bump width in the absence of selective input.

